# A hybrid additive manufacturing platform to create bulk and surface composition gradients on scaffolds for tissue regeneration

**DOI:** 10.1101/2020.06.23.165605

**Authors:** Ravi Sinha, Maria Cámara-Torres, Paolo Scopece, Emanuele Verga Falzacappa, Alessandro Patelli, Lorenzo Moroni, Carlos Mota

## Abstract

Scaffolds with gradients of physico-chemical properties and controlled 3D architectures are crucial for engineering complex tissues. These can be produced using multi-material additive manufacturing (AM) techniques. However, they typically only achieve discrete gradients using separate printheads to vary compositions. Achieving continuous composition gradients, to better mimic tissues, requires material dosing and mixing controls. No such AM solution exists for most biomaterials. Existing AM techniques also cannot selectively modify scaffold surfaces to locally stimulate cell adhesion. We report a hybrid AM solution to cover these needs. On one platform, we combine a novel dosing- and mixing-enabled, dual-material printhead with an atmospheric pressure plasma jet to selectively activate/coat scaffold filaments during manufacturing. We fabricated continuous composition gradients in both 2D hydrogels and 3D thermoplastic scaffolds. We demonstrated an improvement in mechanical properties of continuous gradients compared to discrete gradients in the 3D scaffolds, and the ability to selectively enhance cell adhesion.

## Introduction

Additive manufacturing (AM) techniques offer the automation of the bottom-up manufacturing of 3D structures by adding material selectively where needed. This field has seen tremendous advances in the past three decades, driven primarily by rapid prototyping demands, but gradually being adopted by industries and end users to manufacture unique or small batch products. The techniques have become faster, cheaper, more complex, and better in resolution and quality. It is nowadays possible to build 3D objects by adding material in the form of droplets (1), filaments (2), or to simultaneously crosslink 2D (3) or even 3D patterns (4). The range of materials that can be used today includes metals (5), ceramics (6), polymers (thermoplastics (2), thermosets (7), hydrogels (1, 3, 8)) and several composite materials, such as magnetic particles in polymers (9) and cells suspended in hydrogels (8), depending on the type of AM technology. The field of tissue engineering (TE) and regenerative medicine, which aims to repair or replace damaged tissues in the body, has benefitted greatly from these advances. AM techniques have been gradually re-engineered to accommodate the more demanding requirements of the materials used in TE where cells are seeded into temporary scaffolds to build new tissues (10). Despite the flexibility of some of these new AM techniques, where almost any structure, using any material, for diverse applications seems possible, certain material and application combinations remain surprisingly challenging or even impossible to achieve. One such example is the inability to create constructs mimicking the continuous composition gradients found in multiple tissues and interfaces using most biomaterials (i.e., materials that can be introduced into the body without causing any harm to cells). Continuous composition gradient scaffolds using one of the most common material types for TE scaffolds, viz. biodegradable thermoplastic polymers, remain especially out of reach using existing techniques.

Among biomaterials for TE scaffolds fabrication, thermoplastics offer several advantages for engineering hard musculoskeletal tissues. Their mechanical properties intrinsically resemble the target tissues and can be further matched by tuning the porosity and pore architectures (2); they can be used in simple melt extrusion–based techniques without requiring any solvents; and they can be combined with an inorganic phase, using melt- or solvent-based compounding (mixing) techniques, to resemble bone composition (11). Composition gradients in thermoplastic scaffolds are expected to be advantageous not only for mimicking the composition variations in hard musculoskeletal tissues, thereby enabling improved control over specialized load-bearing abilities and interfacing between tissues, but also for offering the possibility to stimulate cells by varying the physico-chemical properties (12). In the body, highly stiff bone transitions smoothly to relatively softer and more flexible cartilage, tendons or ligaments, and to even more compliant muscle tissue. These stiff and soft juxtapositions help these tissues to bear higher loads than the soft tissues could do alone without large deformation, and dampen better against impacts than the hard tissues could manage alone. Current scaffolds generally lack such transitions.

To generate bone scaffolds harboring these transitions, AM strategies have been previously proposed. Bittner et al. and Gomez et al. have used tricalcium phosphate (TCP)- or hydroxyapatite (HA)-loaded poly(ε-caprolactone) (PCL) and produced AM scaffolds with composition gradients, along the scaffold height or within single layers, by varying the filler concentration using separate printheads for each composition, in a multi-step process (13, 14). Diverse advantages, such as controlling the mechanical properties, degradation rates and spatial distribution of regenerated bone, were cited among the potentials of such gradients. However, the application of separate printheads for each composition limits the number of compositions that can be achieved within a scaffold and the transitions between compositions are stepped and discrete, not continuous.

To obtain the much-needed continuous composition gradients in TE scaffolds, and to achieve them in a single integrated process, we report a printhead design and prototype to enable continuous composition changes during printing using gas-pressure-based ratiometric dosing of two materials and mixing using an extrusion screw. Additionally, the printhead can heat materials up to 250°C and extrude high viscosities, thereby enabling processing of thermoplastic biomaterials with a high filler content. While printheads with ratiometric dosing were reported earlier (9, 15), none so far includes simultaneously high temperature heating and mixing of high viscosity materials. We demonstrate the prototype’s compatibility with high viscosity materials to produce continuous gradients using a range of composite materials developed for bone TE. The produced continuous gradient scaffolds showed improved mechanical properties when compared to discrete gradient scaffolds – with the continuous gradients failing at >50% higher strains on average. Lastly, to test the versatility of the printhead, we performed continuous composition gradient extrusion for hydrogels (lower viscosity than the thermoplastics and room temperature process).

In addition to the printhead prototype, we aimed to include control over the scaffold surface in order to facilitate cell attachment, which is a critical process for tissue regeneration. Plasma activation of scaffold surfaces improves cell attachment. Atmospheric pressure plasma jet (APPJ) systems, which produce a narrow plasma flame, have been integrated on AM platforms to surface-treat the scaffolds during their manufacture (16). With APPJs, uniform activation in deep regions of large, porous scaffolds is possible by applying the treatment during the manufacturing process, as opposed to plasma chambers which are used as a post-processing step and provide limited plasma activation in interior regions of the scaffolds. Additionally, precursor molecules can be used in the APPJ plasma flame to enable the deposition of plasma-polymerized thin films presenting desired functional groups in a mild, solvent-free process.

We incorporated an APPJ system on the same AM platform as the gradient printhead, which allows for a full and uniform treatment of complete scaffolds of all sizes, a potential to mask surface changes introduced by the bulk composition changes in gradient scaffolds, and also offers the possibility to selectively treat regions to promote specific cell attachment areas in a 3D space within the scaffold. The integrated APPJ system was validated by observing the deposition of plasma-polymerized films at desired locations within scaffolds and by observing the adhesion of human mesenchymal stromal cells (hMSCs) on the plasma-patterned regions of the scaffolds.

Combined together, these data demonstrate that our integrated hybrid AM platform offers the ability to create scaffolds with continuous physico-chemical gradients and to selectively activate or coat the scaffold surfaces. This platform offers a wide range of possibilities to push forward the state-of-the-art in TE scaffolds.

## Results

### Printhead design and prototype

A printhead capable of mixing controllable ratios of two thermoplastic polymers during extrusion has been developed and integrated on an AM platform (Fig. 1A, B). The salient design features of the printhead include: (i) two independent material reservoirs feeding a mixing chamber, (ii) an extrusion screw to mix materials, including those with high viscosities, (iii) independently controlled cartridge heaters with integrated thermocouples for the material reservoirs and the mixing chamber heating, (iv) independent digital pressure regulators for the two reservoirs, (v) one-way valves at the interface of reservoirs and mixing chamber to prevent back flows of materials (Supp. Fig. S1A), (vi) dead volume minimization to ensure extrusion of most material that is fed to extrusion screw, (vii) simplified disassembling design to allow the placement of all components in direct contact with the processed materials into solvents for thorough cleaning, and (viii) extrusion needle insulation to prevent its cooling, thereby avoiding the need to overheat material for extrusion, thus indirectly minimizing the chances for polymer thermal degradation. Gradient scaffolds can be printed by simultaneously programming the print path and the magnitude and timing of pressure signals to the two material reservoirs, thereby controlling the composition of the material fed to the extrusion screw. The mixing in the extrusion screw ensures that the composition change is gradual.

**Fig. 1.**
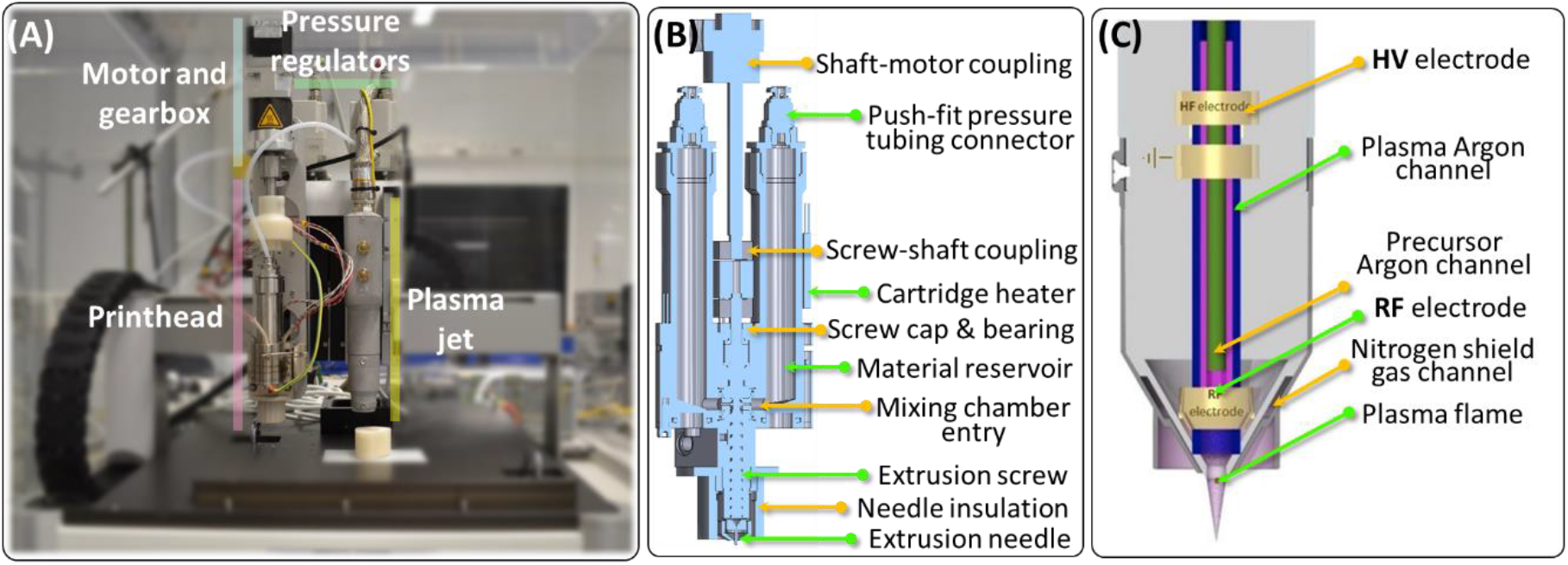
Hybrid AM platform. A printhead for mixing and extruding two materials (**A**, **B**); and an atmospheric plasma jet (**A**, **C**) were developed and mounted on a commercial printer platform (**A**). (**B**) Schematic of the printhead, capable of varying the extruded material composition during the printing, by varying the ratio of the two feed materials using gas pressure applied to each material. (**C**) Schematic of the plasma jet that activates surfaces using an inert gas (Ar tested here) plasma, as well as of depositing films of plasma-polymerized precursors (up to two precursors, fed to the plasma using carrier gas bubbling in the precursors and controlling precursor amount using gas mass flow meters).

### Design of the extrusion screw

The extrusion screw is a crucial part of the printhead, and key design choices were made to ensure mixing capabilities. An extrusion screw, consisting of a helical thread cut into a cylinder, uses rotational motion to transport fluids or granular solids along the axis of rotation while reducing the force necessary to promote extrusion. In addition to this main function of transport, extrusion screws can also provide a degree of mixing resulting from flow patterns. Computational modeling was used to understand flow and mixing behavior in extrusion screws with different thread cuts. Several specialized design features have been reported to modify these flows with the aim of improving mixing (19). Some features divide and re-combine the fluid multiple times to distribute materials being mixed more evenly, while others shear the materials in tight spaces to break aggregates. Both types of features reduce the transport function, and the latter type can additionally increase the torque required to rotate the screw. Besides these tradeoffs, these designs were also too complex to manufacture at the scale required for the printhead, so we selected a design with the thread cut possessing a (filleted) triangular profile (Fig. 2A).The models showed that the triangular cut threads had a smaller dead volume, i.e. volume with low flow velocity, when compared to a square cut of roughly the same cross-sectional area (Fig. 2C). Dead volumes become a source of contamination when the materials are switched in the screw, and therefore should be minimized to reduce cross-contamination while producing the gradient scaffolds.

**Fig. 2.**
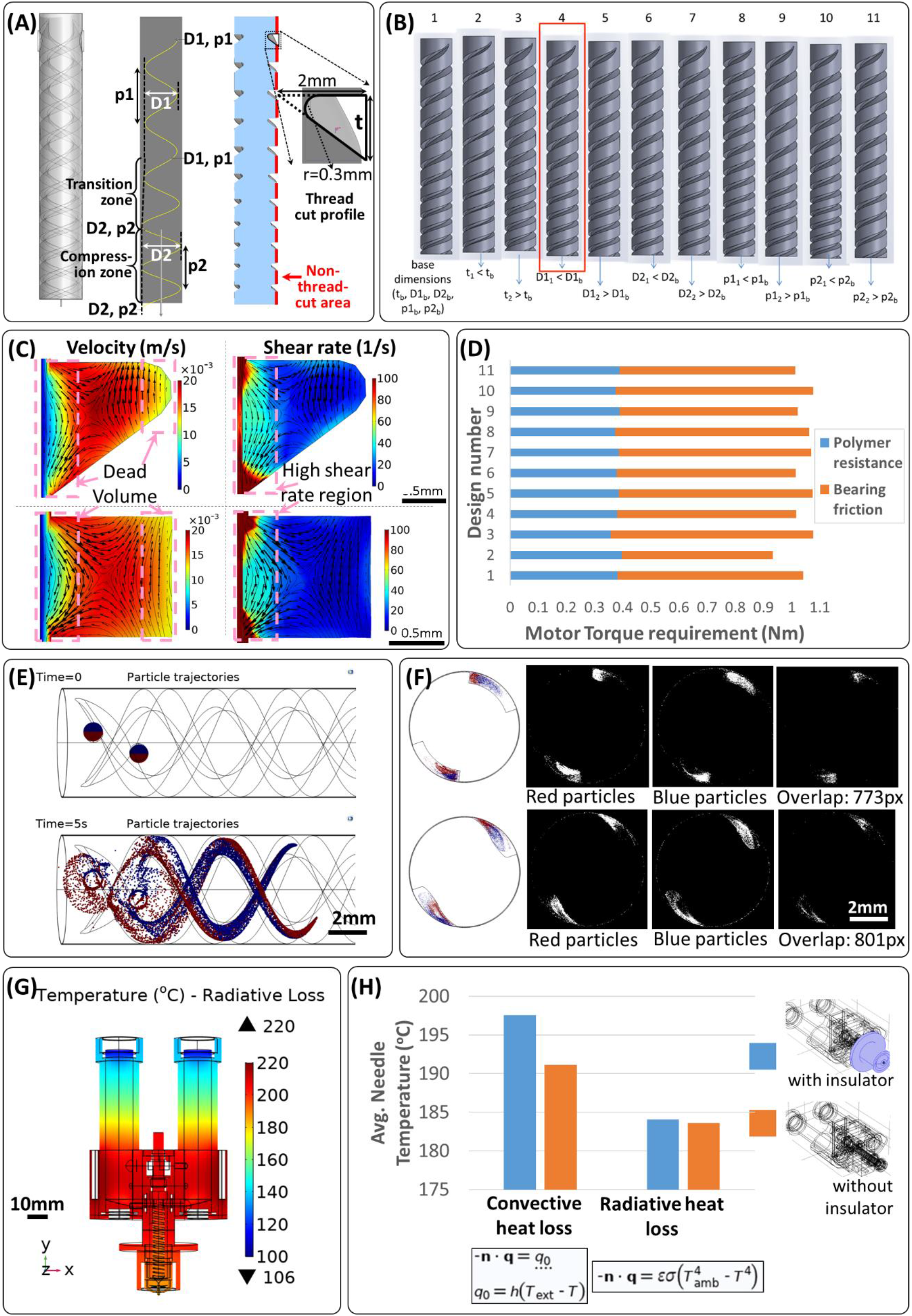
Computational modeling assisted in the screw geometry selection, estimation of torque requirement for the motor, comparison of mixing based on thread cut shape and estimation of thermal insulation provided by an extrusion needle cap. In order to limit printhead weight by limiting the screw operating motor weight, modeling was used to select the screw geometry that reduced the operational torque requirement. The geometrical parameters that could be varied included thread cut width and the depth as well as the pitch of the cuts in the normal and compression zones (**A**). A set of base parameters was selected and variations of each parameter to a higher and a lower value were tested (**B**). In addition to the torque requirement determination, the models provided a way to visualize flow speed and shear rate distributions as well as local flow directionality (**C**) in the various designs tested as well as comparison with a square thread cut profile. Finally, comparing the torque requirements of the design variations tested (**D**) enabled the selection of a screw design for manufacture, design 4 indicated by the red rectangle in panel B. The effect of the screw profile on mixing was tested using a particle tracing model, where transfer of particles starting in one half of the inlet (red) to the region occupied by the other half of the particles (blue) was visualized (**E**). The overlap of the regions occupied by red and blue particles was used as a quantitative measure of mixing (**F**) and to compare a triangular thread cut profile (bottom row) with a square cut profile (top row) of roughly the same area. Lastly, heat flow modeling was used to visualize temperature distributions in the various parts of the printhead (**G**) and compare the utility of extrusion needle insulation against convective and radiative losses (**H**).

Triangular thread profiles have been reported to result in enhanced aggregate breaking by generating a squeezing effect where material gets pulled, the flow being termed as extensional or elongational flow (20). Indeed, the flow streamlines and the magnitude-proportional flow velocity vectors in the plane of thread cross-sections showed the generation of the elongational flows, as indicated by converging flow vectors (Fig. 2C). The flow was mainly along the thread, almost resembling laminar flow, as the 3D velocity vectors were almost parallel to each other and were dominated by the out-of-plane component. Although much smaller than the out-of-plane component, the directionality of the in-plane velocity vectors showed, other than evidence of elongational flows, that the screw motion introduces a gradual twisting of the material inside. Interestingly, streamlines and velocity vectors in the square cut thread also showed converging velocity vectors (Fig. 2C), but due to the larger dead volumes of the square cut, the triangular cut remained preferable. Additionally, the triangular threads provide a larger high-shear-rate region within the thread compared to the square cut, thereby increasing the chance of aggregate breaking (Fig. 2C).

In addition to flow, we evaluated the mixing capacity of the different thread cut profiles using a particle tracing model, which simulates how particles starting in one half of the inlet area mix into the region occupied by the particles starting in the other half as they flow through the screw (Fig. 2E, Movie S1). This mixing model utilized a laminar flow to simplify the visualization and quantification of the mixing. The degree of mixing was quantified as the overlap in the region occupied by one set of particles with the region occupied by the other set. The model showed that the overlap areas were comparable between the triangular and square designs, but slightly higher (801 pixels vs. 773 pixels) for the triangular thread cut (Fig. 2F). While the longitudinal cross-sections used for this evaluation had a larger area for the square thread than the triangular thread (4.6 mm2 vs. 4.17 mm2), the combined area occupied by all the particles was smaller for the square thread (4031 pixels vs. 4612 pixels), as a result of compaction from the larger dead volume in the square thread. When we conducted a mixing model to include the effect of the screw motion for a smaller section of the screw, higher mixing was found for both designs, but with comparable extents between the triangular and the square threads (Movie S1).

After selecting the triangular thread cut shape, the thread cut depth was reduced in the lower section of the screw (Fig. 2A), a common feature known as a compression zone, generally used to homogenize the polymer melt temperature in regions separated from the screw by the poorly heat-conducting polymer and to remove air bubbles by compressing it. A compression zone additionally generates elongational flows as the material is squeezed into a smaller space, contributing to mixing. Because of the triangular shape of the screw thread, reducing the cut depth also reduced the cut width, thus increasing the uncut area on the screw. This was used to increase the total thread length by including more thread coils by reducing the thread pitch, i.e. the distance between the coils (Fig. 2A). This increased the polymer residence time in the screw and hence resulted in better mixing. Finally, two diametrically opposite parallel threads were cut on the screw (Fig. 2A) to give both materials identical access to the screw at any screw position, thereby avoiding the need to synchronize the injection of either material with the screw position.

### Geometric parameters for torque requirements

After the key design parameters for the extrusion screw were selected, we investigated the geometric parameters, including thread cut depth and pitch in the normal and the compression zones, and the thread cut width (as shown in Fig. 2A – t represents the cut width; D1 and D2 the thread cutting tool’s path diameters in the two zones, thus providing inverse measures of cut depth and p1 and p2 being the pitch for each zone). It was important to select geometrical parameters that resulted in the lowest torque requirement, since a smaller motor and gearbox size could be selected reducing the overall size and weight of the printhead. A constant length of the screw (10.25 mm) was always used to linearly transition from D1, p1 to D2, p2. The total length (41 mm) and diameter (6 mm) of the screw were selected based on constraints on total printhead dimensions. Computational modeling was used to simulate the influence of these parameters on the torque required to overcome the viscous (and pressure-based) resistance from the polymer and the normal upwards force exerted on the screw by the polymer as the screw pushes the polymer down. To avoid the transmission of this normal force to the motor and the gearbox, a flange with low friction bearings was introduced at the screw top and bolted to the extruder body. The torque required to overcome the friction between the screw flange and the selected bearing was also taken into account for the selection of the appropriate motor/gearbox solution.

With the generated model for a polymer with 1000 Pa.s viscosity, the total torque requirements to turn the screw varied in the range 0.93–1.08 N.m for the geometrical variations considered. A lower and a higher value of each of the parameters—t, p1, p2, D1 and D2— around a set of base parameters (tb, p1b, p2b, D1b and D2b), was tested (Fig. 2B and Supp. Table S1). Generally, the upwards force on the screw, and hence the bearing frictional component of the torque, was proportional to the volume of polymer in the screw thread and the viscous resistance component was generally dominated by the non-thread-cut area on the screw (shown in red in Fig. 2A), that being the high shear rate region (Fig. 2C). The volume of polymer in the screw was higher when either the thread cut width or the thread cut depth was increased or when the thread pitch was decreased. Increasing the cut width reduced the non-thread-cut area, and due to the triangular cut profile, so did an increase in the cut depth, by increasing the cut width as well. Thus, in two out of three cases, a polymer volume increase in the screw introduced a decrease in the non-thread-cut area. Hence, an increase in the torque requirement to overcome bearing friction was generally accompanied by a reduction in the torque required to overcome the viscous resistance, explaining the overall low difference in operational torque requirements between the designs.

Among the tested conditions, we evaluated the designs offering the lowest torque requirements (Fig. 2D). Design 2 provided the lowest, but the thread cut was very small (Fig. 2B), increasing the risk of filler aggregation that could block the screw when extruding highly filler-loaded materials. Furthermore, manufacturing such a screw with the small thread cut would be challenging, hence this design was not selected. Designs 4, 6, 9 and 11 had the next higher and almost equal overall torque requirements (Fig. 2D). From these, design 4 was selected for manufacture since it allowed the highest compression ratio (ratio between the cut depths in the normal and the compression zones) among those four designs, which would give the highest degree of elongational flow, which contributes to the mixing.

### Computational modeling of extrusion needle insulation

Lastly, computational modeling was used in the printhead design process to evaluate the efficacy of a custom-designed extrusion needle insulation. By using the printhead geometry, the models provided a good understanding of the temperature distributions on the printhead taking into account all the geometrical features (Fig. 2G). Simplified heat transfer models showed that the extrusion needle insulator can be effective against convective cooling, but not against radiative heat loss (Fig. 2H). The actual radiative loss would probably be lower than predicted by the model as the designed insulator was intended to trap air around the needle, which could heat up and lower radiative loss, an effect not captured in the model.

### Validation of the printhead using a rheologically diverse material library

Four polymeric composites, each composed of a distinct filler loaded into a base polymer, were used to validate the printhead. The base polymer was a random block copolymer poly(ethylene oxide terephthalate) / poly(butylene terephthalate (PEOT/PBT), comprising 300 g/mol average molecular weight PEO and a PEOT:PBT weight ratio of 55:45. The filler loadings were: (i) 45% (w/w) HA nanoparticles, hereafter denoted 45nHA; (ii) 10% (w/w) reduced graphene oxide, hereafter denoted 10rGO; (iii) 10% (w/w) zirconium phosphate lamellar filler and 10% (w/w) intercalated gentamicin, hereafter denoted 20ZrP-GTM; and (iv) 10% (w/w) magnesium-aluminum layered double hydroxide and 10% (w/w) intercalated ciprofloxacin, hereafter denoted 20LDH-CFX (21, 22). The first two polymeric composites were developed with the goal of improving the mechanical and bone-formation ability of scaffolds, while the last two were developed to provide local antibiotic release from scaffolds after implantation, avoiding the need for systemic delivery.

Rheological measurements showed that the material library covered a wide range of viscosities (25–40 Pa.s for the polymer alone or with the fillers with antibiotics, ~200 Pa.s for 45nHA, and ~86,000 Pa.s for 10rGO) at empirically determined extrusion temperatures and 1 s-1 shear rate (Fig. 3A). The printhead successfully extruded all the materials, demonstrating its versatility. The high viscosity materials showed large decreases in viscosity at high shear rates, explaining why the extrusion was still possible for highly viscous compositions such as the 10rGO (Fig. 3A).

**Fig. 3.**
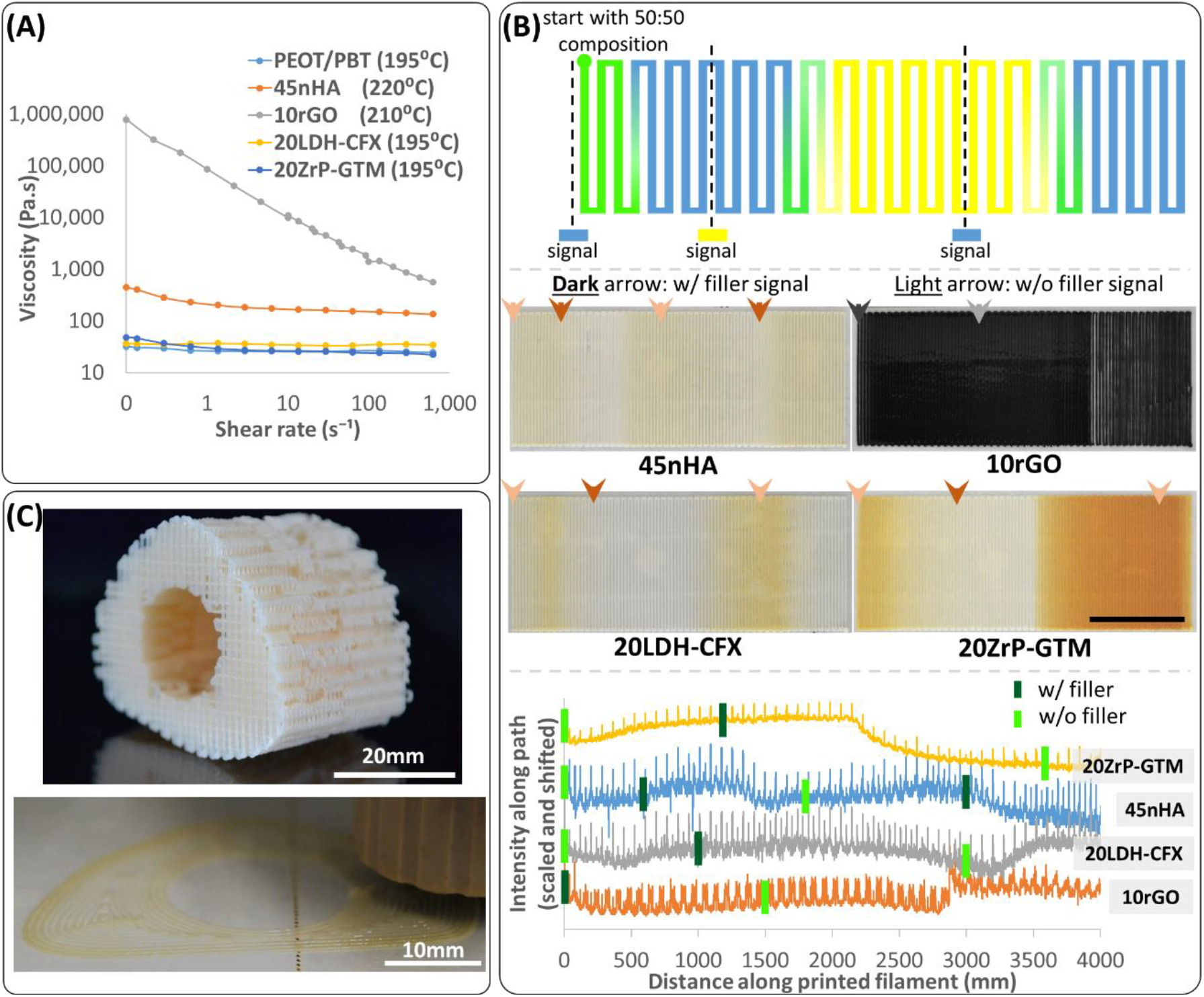
The printhead was validated with a material library developed for bone TE. Extrusion and continuous gradient production were possible with 5 different materials covering a wide range of viscosities. Rheological measurements consisted of viscosity measurements at various shear rates at empirically determined printing temperatures for the materials (**A**). Gradient printing with the selected materials was tested by printing continuously in a single plane while switching pressures from being applied only to one material to only the other material, and monitoring how the extruded composition changed in response (**B**). The top panel in (**B**) schematically shows the printing path using two different materials (shown in blue and yellow), the position along the print path where pressure was applied to one material but not the other (denoted by the signal in the respective color), and how that changed the composition along the print path. The middle panel shows the printed gradients using the four filler-loaded materials switching with PEOT/PBT without filler. The bottom panel shows the intensity of the middle panel images along the print path plotted against the print length. The empirical knowledge of composition switching time after a signal application was used to plan the printing of a long bone defect–shaped scaffold with a polymer-only core and a HA-loaded polymer cortical region (**C**). The top image in (**C**) shows a long bone defect-shaped scaffold printed with PEOT/PBT alone and the bottom image shows a similar scaffold being produced with a HA composition gradient.

### Empirical gradient characterizations and complex gradient scaffolds production

Having determined that the printhead was able to extrude the different polymeric composites, we tested how effectively the printhead could switch material composition to produce complex gradient scaffolds. The time necessary for a complete composition switch between two conditions (base polymer and polymeric composite) was empirically determined for the four combinations. The switching time was calculated by printing continuously a meander in a plane and switching composition from polymer to composite and vice-versa and observing the changes reflected in the extruded filament (Fig. 3B). It was observed that (i) the complete switching generally corresponded to the extrusion of a volume around the screw thread volume (~160 μl, corresponding to a ~1.3 m printed length for 400 μm diameter filaments), (ii) the switch occurs as a smooth continuous composition gradient (instead of a sudden step change), and (iii) introduction of a filler is sometimes more rapid than the complete removal of the filler (e.g., 45nHA and 20ZrP-GTM) (Fig. 3B).

Using the knowledge of the printing length needed to get the desired composition switch, pressure signals were applied during the scaffold printing to get composition changes either within a single layer or at the desired height in a scaffold. For example, human long bone segmental defect-filling scaffolds were printed with a peripheral high and central low HA content (Fig. 3C), resembling a cortical-trabecular mineral distribution in natural bones, where the mineral content gradient is due to a porosity gradient. This printing was done in a single continuous process, starting with printing the central part of the scaffold with PEOT/PBT and switching to 45nHA to print the peripheral region. A layer of another scaffold was then printed, having a central high and peripheral low HA content, using a 45nHA to PEOT/PBT material switch. Printing this second scaffold, together with the printing of even numbered layers of both scaffolds with PEOT/PBT, allowed to start again in the odd numbered layers of the first scaffold with PEOT/PBT in the central region and repeating the composition distribution of the first layer (Movie S2).

In addition to the transition regions between two materials obtained by switching between completely one material to the other, sustained intermediate compositions could also be extruded using a pressure duty-cycle approach. Here, two materials could be fed to the extrusion screw in desired intermediate ratios by varying the ratio of times that pressure is applied to each material in each duty cycle. Applying pressure to only one material at a time ensures that only the one-way valve of that material’s reservoir is open, reducing the risk of material backflow into the other material’s reservoir. To demonstrate this process, we printed a gradual HA gradient scaffold using PEOT/PBT and 45nHA materials, which resulted in a larger number of layers with intermediate HA fraction compositions, compared to the transition region obtained with complete material switching (Movie S3, Supp. Fig. S3).

To further demonstrate the versatility of the printhead, continuous composition gradient production using hydrogels, another biomaterial class useful for bone TE, was also demonstrated. Alginate loaded with HA particles was used for this demonstration, working at room temperature and a relatively low viscosity material (Supp. Fig. S4).

### Mechanical properties of continuous vs. discrete composition gradient scaffolds

In bone TE, mechanically stiffer scaffolds can be manufactured with the addition of inorganic fillers to polymers, which has the drawback of the resulting materials also becoming more brittle for higher volume fractions of filler. In discrete gradient scaffolds, the juxtaposition of stiffer (filler-rich) and less stiff (filler-free) regions is likely to result in high lateral stresses at the interface under compression, due to the mismatch in the mechanical properties. We hypothesized that continuous gradients would reduce these interfacial stresses by reducing the difference in mechanical properties between consecutive layers. In principle, this should also improve the interface between tissues that form in response to stimulation from the two compositions.

We tested our hypothesis by mechanically testing our printed discrete and continuous HA composition gradients in PEOT/PBT scaffolds (Fig. 4A). Two types of gradient scaffolds were tested – (i) 45nHA regions sandwiched between PEOT/PBT regions (PHP) and (ii) PEOT/PBT regions sandwiched between 45nHA regions (HPH). The continuous gradients (PHP_C and HPH_C) were produced by switching during printing between materials (PEOT/PBT or 45nHA). For discrete gradients (PHP_D and HPH_D), the material was switched by pausing the printing, ensuring a complete replacement of the material inside the screw by extrusion away from the scaffold, and then manufacturing the other composition region. Samples with 4 mm diameter and 4 mm (PEOT/PBT, 45nHA, HPH_C; HPH_D), 8 mm (PHP_D) or 7.2 mm (PHP_C) height were cored out from scaffold blocks with a biopsy puncher and used for mechanical testing (Fig. 4A). Three replicates per condition were tested.

**Fig. 4.**
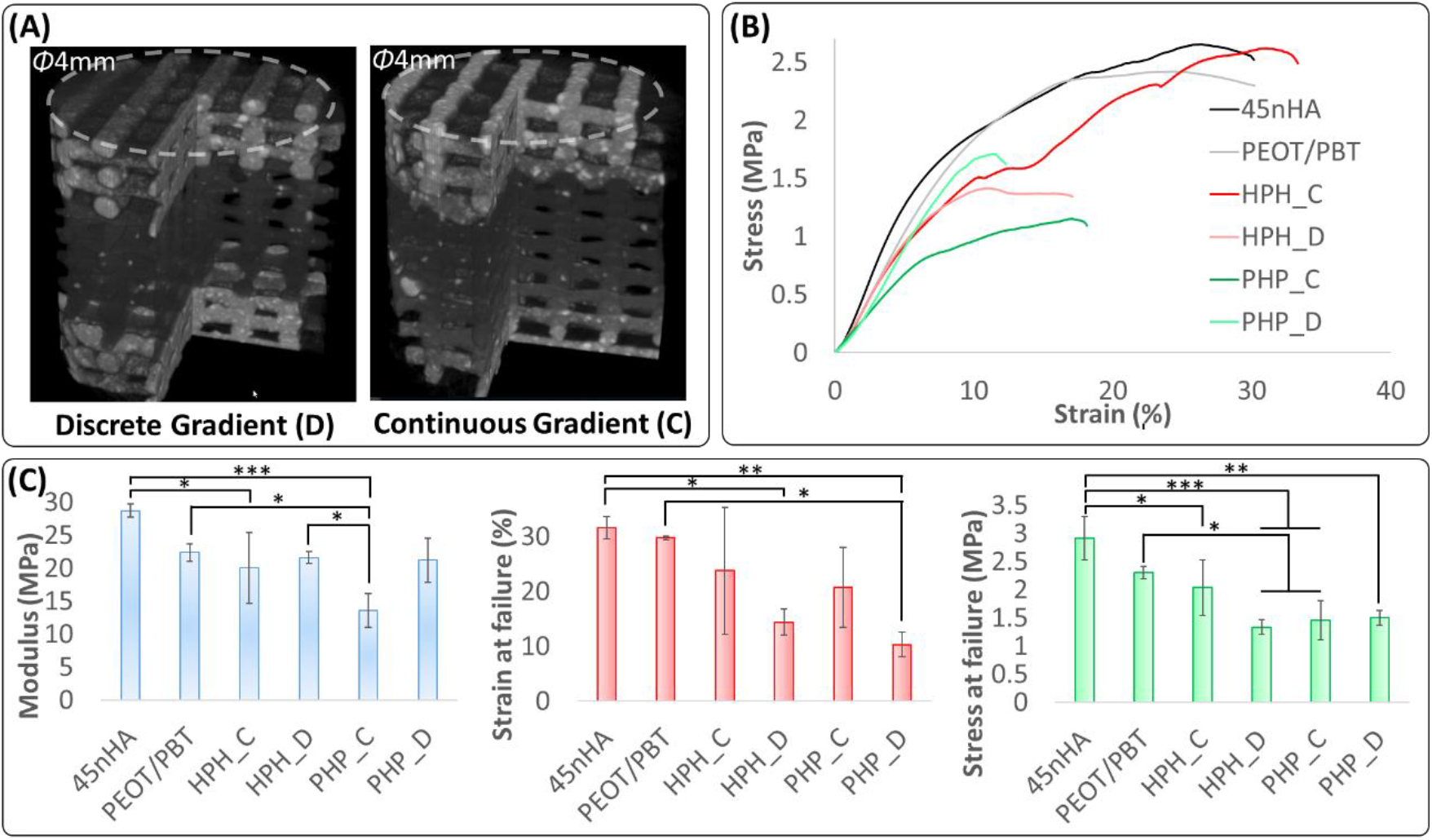
Printing of gradient scaffolds using the developed printhead demonstrated an improvement in the mechanical properties of continuous vs discrete composition gradients. Mechanical test samples were imaged using micro–computed tomography (μCT) to demonstrate the differences in filler distribution between the discrete and continuous gradients (**A**). Uniaxial unconfined compression tests were performed and the comparison of stress-strain curves demonstrated the variations in load-bearing behavior of the various scaffolds (**B**). From the stress-strain curves, the elastic modulus, the strain and stress at failure were computed and demonstrated that continuous gradients withstood higher strains before failure (**C**, plots show mean ± standard deviation for 3 replicates and significant differences are marked as * for p<0.05, ** for p<0.01 and *** for p<0.001).

Mechanical tests of the scaffolds showed that all gradient scaffolds had a similar average Young’s modulus as the control, uniform composition, PEOT/PBT scaffolds (Fig. 4C). The only exception was PHP_C, where the scaffolds showed an appreciably lower average modulus, but the videos (Movie S5) of the tests suggest that this was likely due to parts of the scaffolds shearing and not bearing the load normally, potentially an artefact from shearing during the core out of the test samples from the blocks, which was greater for the PHP scaffolds compared to HPH scaffolds due to a greater scaffold height. Most importantly, the continuous gradient scaffolds consistently withstood higher strains before failing (Fig. 4C and Movies S4, S5) compared to the discrete gradient scaffolds. Failure was taken as the point where stress first dropped by 5% or more from the nearest yield peak. The stress strain curves (Fig. 4B and Movies S4, S5) for the continuous gradient scaffolds generally showed multiple yield points before the failure, while the discrete gradient scaffolds failed after the first yield. In HPH_C, but not in PHP_C scaffolds, the higher failure strains led to higher strengths at failure (Fig. 4C). The lower strength at failure of the PHP_C scaffolds was likely also the effect of the shearing of the scaffolds. Interestingly, the mechanical test videos (Movies S4, S5) reveal that the discrete gradient scaffolds almost always started collapsing at the interface between layers of different material, while in the continuous gradients such was not observed. Thus, continuous gradients demonstrate improved scaffold integrity that is essential for demanding load-bearing applications.

### Design of plasma polymerization–capable APPJ for AM

There are currently no established methods to pattern in 3D the surface of an AM scaffold during the manufacturing process, even though such a capability can be useful to promote cell adhesion in specific regions of interest within the scaffold. A plasma jet integrated on an AM platform provides such an ability. Thus, we combined a commercial atmospheric pressure plasma jet (APPJ, Nadir s.r.l.) (23) in the AM system to allow surface treatment while manufacturing TE scaffolds (Fig. 1A, C, and Fig. 5A). The APPJ has a dielectric barrier discharge (DBD) configuration with the electrodes for plasma generation positioned outside an alumina tube of 6 mm diameter. An alumina capillary is mounted coaxially inside the alumina tube for the supply of chemical precursors. Another tube outside the alumina tube and electrodes provides a confinement atmosphere (Fig. 1C).

**Fig. 5.**
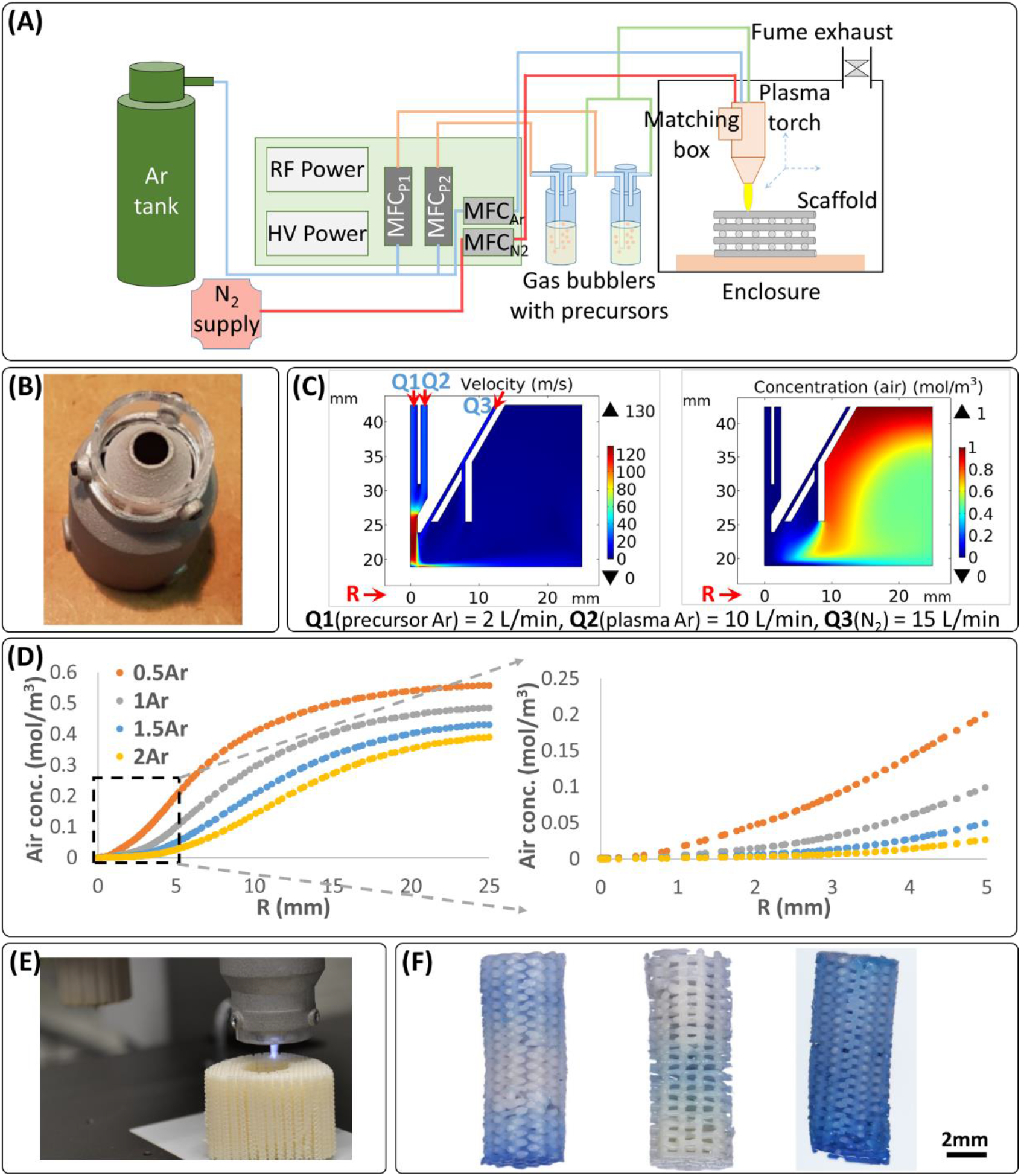
An atmospheric plasma jet was adapted for the AM process, computational modeling assisted in optimizing gas flows for the process, and plasma-assisted patterning of 3D scaffolds was demonstrated. The implemented plasma process consisted of a dual power supply and mass flow controllers for the plasma gas (Ar), shielding/cooling gas (N_2_) and two precursors in carrier gas (Ar) (**A**). The formation of a shielding gas shell around the reactive plasma flame was enabled by a nozzle cap (**B**), and the plasma and shielding gas ratio was optimized with the help of computational modeling (**C**, **D**). A combination of flow and diffusion modeling was used, as depicted by representative flow velocity and diffusion-affected concentration plots (**C**). The models showed that by increasing the flow of the shield gas relative to the plasma gas, a bigger inner volume could be shielded from air, as depicted by the air concentration falling to zero for a larger plasma radius (R), measured near the substrate (**D**). The plasma jet was integrated on the printer platform (**E**), and consecutive printing and plasma treatment layer-by-layer demonstrated the ability to plasma-treat a scaffold fully or at desired locations in 3D, as shown for uniform composition PEOT/PBT scaffolds coated with VTMOS-MAA and stained with methylene blue (**F**). The plasma-treated scaffolds shown in (**F**) are (from left to right) (i) bottom and top 10 layers plasma-treated, middle 20 layers non-treated; (ii) middle 10 layers plasma-treated and top and bottom 15 layers non-treated; and (iii) all layers plasma-treated.

Argon (Ar) was used as the process gas in the alumina tube, in order to reduce the plasma gas temperature at the surfaces being treated, in comparison to molecular gases such as nitrogen (N_2_), which can transfer extra thermal energy to surfaces by relaxing extra rotational states not possessed by Ar. The use of Ar as process gas provides an additionally inert atmosphere for the chemical precursors and avoids their undesired oxidation. Ar is also supplied in the capillary after bubbling through precursor solutions at room temperature. The interaction of the chemical precursors with the high energy electrons of the plasma leads to the creation of radicals and the initiation of chemical reactions. The reaction products then condensate, generating a plasma-polymerized thin coating (thickness in the nanometer range) on the surface of the extruded filaments.

The APPJ works in dual frequency mode by coupling a radio frequency (RF, 27 MHz) power source with a high voltage (HV, 10 kV, 17 kHz) power source. A matching box, based on two manually adjustable capacitors, to reduce the reflected RF power in the plasma, was integrated directly on the plasma jet in order to reduce its weight for the ease of mounting on the printer platform. Both power generators allow working in pulsing mode by selecting a suitable duty cycle, and can also be triggered and synchronized. This particular APPJ design has the key role of obtaining high ion and electron density plasma at low temperature, leading to high deposition efficiency even on dielectric substrates and allowing at the same time to maintain the precursors’ chemical functionalities (24, 25). The RF power source has the main role of controlling the plasma density, while the HV is needed to improve the stability and the plasma interaction with the substrate and to avoid the transition to RF γ-mode, where localized effects start to dominate and form hot filamentous plasma instead of a cold uniform plasma flame (26). N_2_ was used as a cooling and a shielding gas and confined around the plasma flame by the addition of a nozzle cap (Fig. 5B).

### Computational modeling for the plasma process

An important requirement for the plasma jet is the shielding of the reaction volume from atmospheric oxygen, which could quench the free radical polymerization. Computational modeling was performed to investigate the influence of the gas flow rates on this shielding effect. The shield gas (N_2_) flow required was determined using a diffusion model solving for the amount of air diffused into the plasma flame and shield gas volumes at steady state. Since gas flow velocities and pressure were expected to show large spatial variations within the volume of interest, a multiphysics model combining flow and diffusion was used, where the flows calculated in the flow model were used as convection velocity field input for the diffusion model. Due to the tight confinement of the gases within the plasma torch and the low characteristic length (< 1 cm) of the region of interest outside the plasma torch, Stokes flow was assumed and used for the model. Effects of temperature increase from the plasma flame, higher concentrations than assumed (1 mol/m3 was assumed for all gases) and other flow disturbances were bundled as a two orders of magnitude factor of safety in the diffusion coefficients (10-3 m2/s assumed for all gases, 10-5 m2/s is a realistic value for most gas pairs around atmospheric pressure and room temperature (27)). This assumption made the model demonstrate air penetration into the reaction volume, similar to what happens in the real scenario, albeit due to other factors. We were then able to investigate the role of the shell gas in blocking this penetration. The models showed that increasing the shield gas volumetric flow rate more than 1.5 times above the plasma gas volumetric flow rate did not appreciably increase the thickness of the shielded region (determined by the distance from the center of the flame where diffused air concentration was close to 0) (Fig. 5C, D). Thus, a shield gas flow rate 1.5 times the plasma gas flow rate was used for the experiments and showed successful deposition of plasma-polymerized films.

### Plasma-patterned and selective 3D cell adhesion in 3D scaffolds

The APPJ was integrated on a printer platform (Gesim Bioscaffolder 3.0, Fig. 1A) and used to treat defined areas within the 3D scaffolds during the AM extrusion process (Fig. 5E). The ability to feed two precursors to the plasma flame simultaneously enabled the scaffold functionalization with carboxylic groups using a mixture of vinyltrimethoxysilane (VTMOS) and maleic anhydride (MAA). The coated regions could be visualized using non-specific electrostatic adsorption of methylene blue to the coating (Fig. 5F and Fig. 6B).

**Fig. 6.**
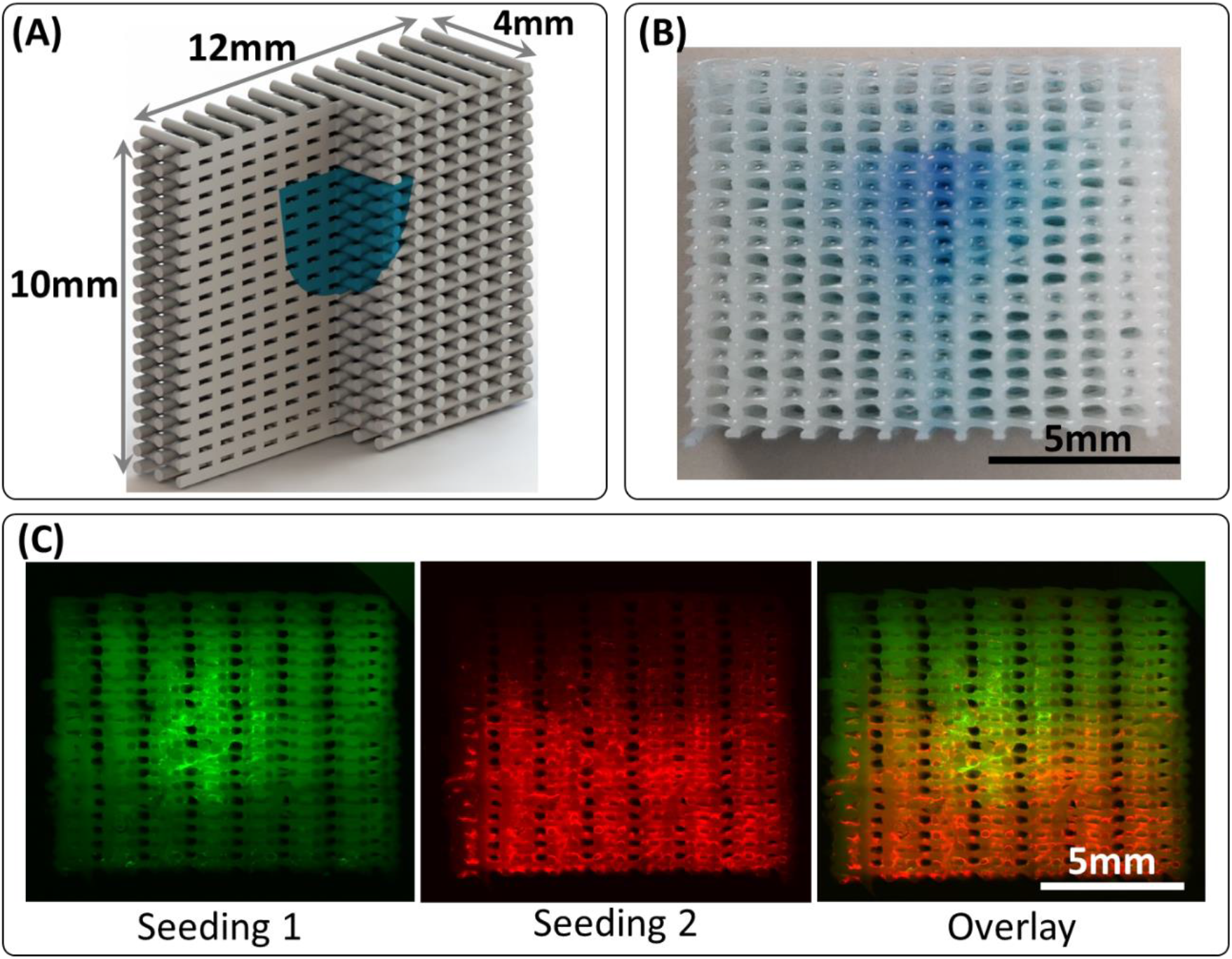
Plasma treatment of scaffolds demonstrated successful patterning of cells deep within a 3D scaffold. A plasma-treated island was patterned deep within a scaffold, as depicted schematically, with the plasma-treated region colored blue (**A**). The patterned region was visualized for the plasma-polymerized VTMOS-MAA coating using methylene blue staining (**B**). The plasma-treated island was used to attach hMSCs stained green selectively, shown here for an Ar-only plasma-patterned scaffold (**C**, Seeding 1). A second cell seeding using a viscous culture media (28) was then used to attach hMSCs stained red on the rest of the scaffold (**C**, Seeding 2), demonstrating an approach to pattern two cell populations (overlay) using the plasma treatment.

To demonstrate that such a selective treatment could be used to pattern cell populations, dual cell seeding was performed on printed PEOT/PBT uniform composition scaffolds treated with APPJ in a defined internal region (Fig. 6A, B). Human mesenchymal stromal cells (hMSCs) were seeded on these scaffolds, and they selectively attached to the plasma-treated region but not untreated PEOT/PBT regions, a typical behavior for this untreated polymer (Fig. 6C). To add a second population of cells to the untreated zones of the scaffold, a second seeding step was done with hMSCs re-suspended using a culture medium viscosity modifier that ensured a slower cellular sedimentation and allowing more time for cells to attach throughout the scaffold irrespective of the plasma treatment (28) (Fig. 6C). With this approach a selective population of cells could be attached to the plasma-treated areas and a second populations of cells could be dispersed throughout the scaffold, showing the patterning potential by controlling the cell adhesion behavior with plasma. Since the first seeding did not cover the entire plasma-treated surface area, cells from the second seeding could also attach to the free available space in the plasma-treated region as well as throughout the scaffold.

## Discussion

We report here advances in AM technologies that address materials and applications not addressable by any other techniques. We developed a printhead capable of creating continuous gradients by dosing desired ratios and mixing of high viscosity materials at high temperature during the printing process (Fig. 1A, B). Such a solution was not available previously, and we also demonstrated its high flexibility to create gradients with low temperature, low viscosity materials, such as hydrogels (Fig. S4). The printhead reduces an otherwise multi-printhead, multi-step procedure to an integrated, single-step process, which saves time and effort. Importantly, the developed printhead allows the production of both discrete gradient and continuous gradient scaffolds with smooth composition transitions. The printhead was validated using a rheologically diverse material library, and the printed PEOT/PBT scaffolds with a continuous gradient in the HA-loading fraction had superior mechanical performance when compared to a PEOT/PBT–HA scaffold with discrete gradients (Fig. 4).

In addition, we adapted an existing APPJ and integrated it on an AM platform to provide a hybrid AM printing and plasma-treatment process. The unique ability of the printer-mounted plasma jet was demonstrated by patterning a cell-adhesive island deep within a scaffold and using it to selectively adhere a cell population, which would be difficult to achieve in a completely fabricated scaffold. The implementation of the plasma polymerization process and the demonstration of cell-patterning using selective plasma treatment on a 3D scaffold are a significant advance beyond the previously reported combination of a plasma with AM process, where no polymerization or patterning was demonstrated (16). Additionally, the previously reported plasma system used N_2_ as a working gas, which produces a higher surface temperature and hence greater surface defect susceptibility than Ar, which we used. Plasma polymerization offers a plethora of features that are unachievable by other existing techniques applied to TE scaffolds. For example, it can be used to leave selected areas untreated and hence non-cell-adhesive, offering a way to guide cell distribution on scaffolds in order to improve tissue regeneration (17). Plasma treatment patterning could also be used to achieve islands of different cell populations, which can be useful for TE strategies requiring multiple cell types. For example, the vascularization of a scaffold could potentially be improved by patterning an endothelial cell 3D core to attract angiogenic progenitors (18).

The developed hybrid platform opens endless possibilities to create different gradient and non-gradient scaffolds with the possibility to fine-tune material composition and surface properties that can be valuable assets for the next generation of thermoplastic scaffolds for a plethora of TE applications. Cell responses in different fabricated scaffolds—such as continuous vs. discrete composition gradients, shallow vs. steep continuous composition gradients—remain to be tested. Additionally, combinations of diverse material types may be explored. For example, materials with varying swelling behavior can be combined to program deformation behavior in the produced scaffolds (9). It would be interesting to see how far the mechanical integrity improvements can be pushed beyond simple composition juxtapositions by utilizing continuous gradients to combine materials that do not adhere well to each other or possess large differences in mechanical properties or both. A very useful test of utility of our hybrid platform would be to determine whether surface coating with plasma-polymerized films could influence cell adhesion behavior to ignore signals such as changes in wettability and surface chemistry, during composition changes in the scaffold. Patterning a plasma-polymerized film-coated island at the composition change interface (Movie S6 and Supp. Fig. S5) could allow for the simultaneous testing of cell response to the two compositions and the interface in absence as well as presence of the plasma polymerized film coating. Thus, we conclude that many exciting possibilities lie ahead using the developed hybrid platform technology.

## Materials and Methods

### Technical details of the printhead and its implementation

The screw and jacket were manufactured using tungsten carbide for high wear resistance, allowing for the processing of polymers loaded with potentially abrasive ceramic particles. The screw thread was defined by the cut profile instead of the thread profile to simplify the manufacturing by reducing the amount of material to be cut out and defining the tool shape.

The materials to be extruded can be loaded into the printhead in the form of pellet, powder or any other format that fits in the reservoirs. Teflon plungers were used in the reservoirs, above the material (Supp. Fig. S1B), to ensure a uniform compression of the material towards the screw while minimizing residual material attached to reservoir walls. The extrusion screw is operated using a stepper motor with a gearbox controlled by a Trinamic controller board and software. The pressure regulators (Supp. Fig. S1C) and the cartridge heaters are controlled using a custom-written LabView program (Supp. Fig. S2B) communicating via a NI Compaq DAQ system coupled with solid-state relays (Supp. Fig. S1D). These printer-independent controllers allow for the easy integration of the printhead on diverse printer platforms, achievable using only mechanical coupling and two electrical triggers from the printer programmable logic controller (PLC) run through the printing G-code: one to actuate the stepper motor and one to actuate the pressures. The printhead was developed within the weight and volume constraints of a commercial printer platform (Gesim Bioscaffolder 3.0) (Fig. 1A), and was integrated using a mechanical coupling, software adaptation, and the two separate triggers (0 to 24 V step changes to designated ports on the PLC).

The printer software was modified to support the integration in the following ways: (i) the motion range of the Z-drive on which the printhead was mounted was programmed for the safe movement of the printhead, (ii) a user interface (Supp. Fig. S2A) was provided for entering printing parameters to automatically create G-codes defining the movements of the printhead and the plasma jet as well as the appropriate triggering of the screw motor and the plasma jet, and (iii) the possibility to add triggers for pressure switches and stepper motor actuation through G-code customization. The Trinamic software was used to control the motor, allowing to tune settings such as the speed and rotation direction of the motor. The triggers from the G-code then switched the motor on/off to be operated at these defined settings. Finally, the LabView program allows for the following: (i) setting, real-time monitoring, plotting and logging of the temperatures applied to the two reservoirs and mixing chamber, (ii) manually applying desired pressure to each pressure regulator (Manual Control button), (iii) applying desired pressure profiles with desired phase lag to each pressure regulator (sine, square, triangle and constant profiles possible) and switching to another pressure profile for each regulator on each pressure trigger from the G-code (Automatic Control button), and (iv) uploading custom-written pressure profiles (each profile an array of pressure values for each regulator to be applied after every 0.1 s), such that each time the G-code triggers the pressure, a new profile could be applied (Raw Control Button) (Supp. Fig. S2B).

### Computational models for screw design selection

COMSOL *Rotating Machinery, Laminar Flow* models were solved for the various screw designs to determine the flow profiles and the resulting resistance to rotation and the upward normal force applied to the screw. Static (*Frozen Rotor*) models were used in the calculations that approximate the screw motion effects by accounting for Coriolis and centrifugal forces resulting from the screw motion. The geometry used was the fluid volume between the screw and the barrel (cylinder at 100 μm separation from the screw), in the inlets (diametrically opposite 3 mm diameter lateral cylinders near the top of the screw) and in the extrusion needle (200 μm diameter, 1 mm long cylinder) (Fig. 2A). A cylindrical surface at 50 μm separation from the screw (halfway between screw and barrel) was used to divide the total volume into a rotating and static (*rotationally invariant*) domains, which was a simplification implemented by COMSOL to avoid moving meshes in a time-dependent solution. The fluid properties used were a 1000 Pa.s viscosity and 1000 kg/m^3^ density. The boundary conditions used were 1 MPa (10 bar) pressure at the inlets, no slip condition at all walls, flow continuity at the domain dividing surface, and 0 Pa pressure at the extrusion needle outlet. The screw rotation speed was fixed at 60 rpm. Once the models were solved, the torque vector directed along the screw rotation axis (z-axis) was calculated as the surface integral over the entire screw surface of the cross product of the *Total Stress* (accounts for both viscous and pressure forces) vector components in the plane perpendicular to the rotational axis (components in the xy plane, (rmspf.T_stressx, rmspf.T_stressy)) and the direction vector from the rotation axis (x and y coordinates (x,y) since axis was centered at (x,y)=(0,0)) for all points on the surface, given by the expression:

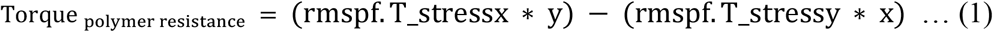

The normal force on the screw was calculated as the surface integral over the screw surface of the component of the *Total Stress* vector along the rotational axis direction (expression integrated: rmspf.T_stressz). The normal force (*N*) was assumed to be equally distributed on the screw flange (3 mm (*R_1_*) to 5.5 mm (*R_2_*) annulus) and friction coefficient (*μ*) of 0.4 (from bearing specifications) was used to calculate the torque as:

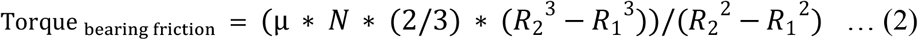

Two geometrical parameters used in the model differed from the actual printhead geometry: the screw-barrel gap was 20 times higher than the real geometry (simplification implemented to prevent an extremely dense mesh in the gap, which would increase the computational cost of the model) and the inlets were bigger than in the manufactured prototype. The inlet size did not affect the viscous drag on the screw so the expected impact on the model was minimal. The screw-barrel gap simplification led to an order of magnitude underestimation of the torque. However, an order of magnitude higher torque was not the correct requirement either, since high viscosity materials that were used with the printhead had lower viscosities at high shear rates. This effect could be included into the models by changing the viscosity from a constant value to a *Carreau-Yasuda* model. This was done for a representative model with the correct screw-barrel gap and reasonably assumed *Carreau-Yasuda* parameters, and showed that the normal force on the screw was similar irrespective of the screw-barrel gap and viscosity model used, but the viscous resistance did increase roughly an order of magnitude for a constant viscosity, and dropped to roughly twice the value from the simplified gap models when the shear rate-dependent changes in viscosity were accounted for (data not shown).

### Computational model for degree of mixing in screw

For the mixing models to check the effect of the screw thread profile alone on mixing, models excluding the effect of the screw motion were used. A COMSOL *Laminar Flow* model was used to only calculate the flow under a pressure difference (0.6 MPa at each of the inlets and 0 Pa at the outlet). A *Stationary* solution was calculated. The geometry was similar to the torque determination models, with a few exceptions. The volume at the base of the screw and the extrusion needle were removed in order to speed up the flows in the absence of the screw motion, the inlets were positioned as in the manufactured printhead (1 mm diameter inlets, diametrically opposite, but vertically de-phased by 2 mm center-to-center, to avoid seeing the same thread at any point during the screw motion), and screw-barrel separation was reduced to 25 μm. The fluid viscosity used was 35 Pa.s (PEOT/PBT viscosity at printing temperature) and density used was 1000 kg/m^3^. After solving for the flow, a COMSOL *Particle Tracing* model was solved that used the calculated flows to drive the motion of 10 μm, 2200 kg/m^3^ particles experiencing a *Drag Force* from the fluid, but not affecting the flow or other particles. Uniformly distributed 10,000 particles were released at each inlet at time 0, and a time-dependent study was solved to track their movement until the 300 s time point. The positions where the particles crossed the bottom of the screw were marked as a 1 pixel × 1 pixel point (corresponding to 20 μm × 20 μm) on a 400 pixel × 400 pixel image representing a 8 mm × 8mm area centered around the screw. This was done separately for particles starting in two halves of the inlet. A binary intersection of the images was obtained for the particles starting on separate sides of the inlet, thus provided an estimate of how many particles from one side had gone over to the other side during the flow through the screw thread, to investigate the mixing effect of the two screw thread cuts investigated (square and triangular).

To include the effect of screw motion in the particle tracing based mixing models, without making the models extremely computationally heavy, only the top 11 mm section of the screw was used. The same geometry and model used for the torque determination were used, except that the inlets were sized and positioned to release full contents to a screw thread at the start, and screw motion was solved for using a time-dependent study. This model included the actual changes in the screw position instead of approximating their effects by the addition of Coriolis and centrifugal forces. The particle tracing conditions used were the same as those for the laminar flow mixing model described above, except that the *Drag Force* was applied to particles only in the *Rotating Domain*, which made the models solve faster without noticeably affecting the flow patterns of the particles. The *Rotating Machinery, Laminar Flow* and the *Particle Tracing* physics were simultaneously solved for in a time-dependent study.

### Computational model for extrusion needle insulation

A COMSOL *Heat Transfer in Solids* model was used to simulate heat loss on the extrusion needle region of the printhead. The full printhead geometry, excluding the cartridge heaters, the motor and the motor shaft, was used and the hollow volumes were filled with a solid. The parts composing the printhead were defined with the material properties from the COMSOL material library: 316L steel for the printhead body, tungsten carbide for the screw and the barrel around it, PEEK to insulators and polyetherimide as a sample polymer to the solid filling the hollow volumes. The walls of the four cartridge heaters were defined at a fixed temperature of 220 °C and the surroundings were defined at a temperature of 20 °C (simulating room temperature standard operating condition). The PEEK parts surfaces were either set as *Thermal Insulation*, i.e. no heat loss boundaries or suppressed, when models without extrusion needle insulation were solved. The remaining surfaces were given a *Surface-to-Ambient Radiation* boundary condition with emissivity 0.5 to model radiative losses or a *Convective Heat Flux* with the *External Natural Convection, Vertical Thin Cylinder (diameter 5 cm, height 15 cm)* heat transfer coefficient to model convective losses. *Stationary* solutions were computed for all cases and the surface average of the extrusion needle temperature was evaluated.

### Rheology measurements

Rheological measurements were made using a TA Discovery HR-1 rheometer under a N_2_ atmosphere to prevent thermal degradation of the polymer. Measurements were made using 25 mm diameter steel parallel plates. The plates were brought to the desired temperature and the plate separation was calibrated. Then, sufficient material was placed on the bottom plate to form a >1 mm thick melt sheet covering the entire plate. The top plate was then lowered until the molten polymer or composites were evenly distributed throughout the plate surfaces (final gap between 500–1000 μm), and the excess material was scraped away. This was done as fast as possible and the environment control chamber was closed. Once the temperature stabilized at the set temperature, a 5 min time sweep was carried out to measure the viscosity in replicate (n=~10), but also to homogenize the melt temperature. The frequency sweep was then carried out in the shear rate range 0.1 s^−1^ to the maximum possible (generally in the range 600–1000 s^−1^).

### Printing parameters for empirical switching time determination

The printing parameters were empirically optimized with the aim of achieving at least one transition from one material to the other material. In all cases printing was started with a 50:50 mixture of the materials contained in the two reservoirs. This was done to save material because at the start, uncontrolled amounts of both materials flowed into the screw, and going to 50:50 composition was easier (by extruding for some time with equal pressure applied to both reservoirs) than clearing the mixing chamber with one of the materials. For printing, the PEOT/PBT reservoir was kept at 195 °C. The remaining printing parameters are summarized in Supp. Table S2. The G-code was generated to produce a continuous meander covering an area of 40 mm × 100 mm with 40 mm long parallel lines spaced 1 mm apart. Pressure switch positions were determined jointly by the printing speed and the pressure profile frequency.

### Printing parameters for radial gradient segmental bone defect scaffold

The G-code was generated from a .stl file using Slic3r. Separate G-codes were created using concentric and rectilinear infills and were combined to print with concentric and rectilinear infills in alternate layers. Printing of a second scaffold and the inclusion of the pressure triggers and motor actuation were implemented by modifying the G-code. The printing sequence used was: fill screw with PEOT/PBT, apply pressure only to 45nHA and start printing the first layer (concentric infill), printing length ensures that extruded material changes to 45nHA in the cortical region, switch pressure to only PEOT/PBT and print a second scaffold on the side (inverted gradient), print second layer of both scaffolds (rectilinear infill) with PEOT/PBT and repeat the process from the 3^rd^ layer onwards. The pressure switches included in the G-code ensured a timely trigger at the appropriate points to switch between constant pressure profiles within the Automatic Control functionality of the pressure control software. The printing temperatures used were: 210 °C for the 45nHA reservoir, 195 °C for the PEOT/PBT reservoir, and 220 °C for the mixing chamber. A 400 μm internal diameter extrusion needle was used and the screw speed used was set at 60 rpm. The printhead translation speed used was 30 mm/s and the pressures applied to each reservoir were 0.87 MPa for both materials.

To better visualize the composition changes, an identical scaffold was printed using dyed PEOT/PBT instead of 45nHA (Movie S2). The print parameters used were the same as for 45nHA, except that the dyed PEOT/PBT reservoir was heated to 195 °C.

### Printing parameters for HA gradient scaffold with slow composition change

The G-code was generated using the printer software interface and the scaffold was printed with a 400 μm inner diameter extrusion needle. The scaffold parameters used to create the G-code were: circumradius of the square scaffold base 15 mm, total scaffold height 10 mm, filament spacing 1 mm, layer height 0.32 mm, printhead translation speed 15 mm/s. The screw speed used was 30 rpm. The pressure trigger signals were inserted in the G-code at the start of layers 1, 8, 15 and 22 (total 31 layers) and the triggers were used to activate pressure profiles uploaded in the Raw Control part of the pressure control software. Printing was started with the screw filled completely with PEOT/PBT. The pressure profiles were prepared to apply the following order of signals at each trigger: 25% 45nHA, 50% 45nHA, 75% 45nHA, 100% 45nHA. This was achieved by applying alternating pressures to the two materials every 4 s, where for the 25% 45nHA profile, the 45nHA would be pushed for 1s and the PEOT/PBT would be pushed for the next 3s, and so on. The pressure applied to either material was kept constant at 0.87 MPa.

### Printing parameters for hydrogel composition gradient demonstration

The materials used were 6% (w/w) low (unknown) molecular weight alginate (Sigma-Aldrich) in water and 16.7% (w/w) hydroxyapatite nanoparticles (nHA) in 6% (w/w) alginate. The G-code used was the same as that used for the empirical material switching time determinations. The screw speed used was 15 rpm and the printhead translation speed used was 6 mm/s. The starting composition was not controlled and pressures were manually switched based on visual observation of composition changes. The pressures applied were 10.8 kPa to the alginate alone and 36 kPa to the alginate containing nHA. The extruded material was deposited on 1 M calcium chloride-soaked tissue paper, to crosslink the alginate as it was deposited. Printing was also repeated using a modified G-code, which increased the separation between filaments to 2 mm from 1 mm to reduce fusion of filaments.

### Printing parameters for continuous and discrete gradient scaffolds for mechanical testing

All G-codes were generated using the printer software interface. For the 45nHA, PEOT/PBT, 45nHA-PEOT/PBT-45nHA continuous (HPH_C) and 45nHA-PEOT/PBT-45nHA discrete (HPH_D) samples, square base scaffolds with circumradius 20 mm and height 4 mm were printed and the mechanical test samples were cored out using a 4 mm diameter biopsy punch.

For the PEOT/PBT-45nHA-PEOT/PBT continuous (PHP_C) and PEOT/PBT-45nHA-PEOT/PBT discrete (PHP_D) samples, square base scaffolds with circumradius 10 mm and height 8 mm were printed and the test samples were also punched out using a 4 mm diameter biopsy punch. The printing parameters for the various scaffolds are summarized in Supp. Table S2.

### Scanning electron microscopy

Micrographs were acquired using a FEI XL30 scanning electron microscope operating in the back-scattered electron mode. Images were acquired using a 25 kV electron beam with spot size 4.

### Micro-computed tomography

Images were acquired using a Scanco Micro-CT100 machine at a 70 kV voltage, 200 μA current, 300 ms exposure and 10 μm isotropic voxel size.

### Mechanical testing and analysis

Compression tests were carried out on a TA ElectroForce 3200 setup using a 50 N or a 450 N load cell, 10 mm diameter compression plates, and strain controlled ramped loading at 0.5% strain/s. Three replicates per condition were tested. Moduli were calculated as the slope of the linear trendlines fitted to the stress-strain data between 2 and 4% strains, since all plots were linear in that range. Failure points were found using a custom MATLAB script to find the local maxima in the stress strain plots and then locating the first place where the stress dropped by 5% after a local maxima.

### Computational model for plasma gas flow and oxygen/air diffusion

A 2D axisymmetric geometry was set up in COMSOL for the plasma jet nozzle along with the nozzle cap forming the shield gas shell around the plasma reaction volume. The geometry also included a substrate at a 5 mm separation from the nozzle and a sufficiently large gas volume surrounding the setup. The models used a 2 mm diameter plasma nozzle, although the manufactured torches often are given a larger diameter to reduce the gas flow velocities. Materials were appropriately assigned to the various parts from the COMSOL material library. First, a *Stokes Flow* model was solved to determine the velocity fields. *Compressible Flow* with a *RANS k-ε* turbulence model was used. The temperature used was 20 °C. The entire gas volume was given the properties (viscosity and density) of Ar and flow rates for the plasma Ar, precursor Ar and shielding N_2_ were applied as inlet boundary conditions at the appropriate boundaries in the geometry. All solid walls were given a 3.2 μm *Sand Roughness* as a way to apply no-slip condition in the turbulence model. The inlet velocities were adjusted to effectively apply a 10 L/min volumetric flow to plasma Ar and 2 L/min volumetric flow to precursor Ar. The flow of the shield gas was varied (5, 10, 15 and 20 L/min) and its effect was studied. Next, a COMSOL *Transport of Diluted Species* model was solved using the flow fields from the *Stokes Flow* model as the convective flow field input. At each of the 3 gas inlets, the diffusing species was given a concentration of 1 mol/m^3^ and all species were given a diffusion coefficient of 10^−3^ m^2^/s. The remaining gas boundary at the top (air region) was also given an inlet boundary condition of 1 mol/m^3^ diffusing species concentration and 10^−3^ m^2^/s diffusion coefficient. In both models, the remaining gas boundaries were defined as 0 Pa pressure outlets. *Stationary* solutions were evaluated but the flows were ramped up in an *Auxillary Sweep* from plasma Ar flow rate starting at 0.5 L/min and increasing in 0.5 L/min steps and the other flows maintaining their respective ratios to it. Finally, the concentration distribution of the air species was evaluated along a line parallel to the substrate and 100 μm above it.

### Printing and plasma treatment parameters for layer-by-layer treated and patterned scaffolds

The initial scaffolds to demonstrate layer-by-layer treatment and 3D patterning ability (Fig. 5F) using the plasma jet were produced using PEOT/PBT, heated to 195 °C in the reservoir and 205 °C in the mixing chamber, pushed using 0.88 MPa pressure and extruded through a 340 μm inner diameter extrusion needle. For each condition, a 5 mm circumradius square base and 10 mm height scaffold block was printed with center-to-center filament spacing of 850 μm and layer height of 250 μm. The screw speed used was 30 rpm and the printhead translation speed used was 17 mm/s. Three types of plasma-treated scaffolds were produced: (i) bottom and top 10 layers plasma-treated, middle 20 layers non-treated; (ii) middle 10 layers plasma-treated and top and bottom 15 layers non-treated; and (iii) all layers plasma-treated. The plasma parameters used were: N_2_ flow rate, 7.5 L/min; plasma Ar flow rate, 5 L/min; RF power, 20 W; pulsing, 250 μs ON / 1000 μs OFF; VTMOS Ar flow rate, 124 ml/min; MAA Ar flow rate, 1.875 L/min; plasma jet translation speed, 17 mm/s; and plasma nozzle to scaffold top surface separation, ~6 mm during each treatment.

For the cell island patterning experiment (Fig. 6A, B), the G-code was created to print a 12 mm × 4 mm × 10 mm scaffold using a 340 μm inner diameter extrusion needle. The scaffold parameters used were: filament spacing, 850 μm; layer height, 250 μm; printhead translation speed, 15 mm/s. Screw speed used was 25 rpm, and the reservoir was heated to 195 °C and the mixing chamber to 220 °C. The pressure applied was 0.72 MPa. When the scaffold was printed to 7.5 mm height, the plasma treatment was carried out, the printing was resumed until a 10 mm scaffold height was achieved. The plasma parameters used were: plasma Ar flow rate, 10 L/min; N_2_ flow rate, 15 L/min; precursors Ar flow rates: 0 for plasma activation only, 0.233 L/min (VTMOS) + 1.764 L/min (MAA) for the plasma-polymerized coating; plasma power, 15 W; no plasma pulsing; plasma nozzle to scaffold top separation, 6.5 mm and plasma translation speed, 1 mm/s. When precursors were used, the flows were allowed to run for 60 s with the plasma positioned away from the scaffold to allow for the precursors to reach the flame.

For the demonstration of the combination of bulk composition gradient with plasma patterning, the same settings were used. The only difference was that the printing was started with only PEOT/PBT in the screw and pressure applied to dyed PEOT/PBT.

### Cell culture and imaging on plasma-patterned scaffolds

HMSCs isolated from bone marrow (Texas A&M Health Science Center, College of Medicine, Institute for Regenerative Medicine, donor d8011L, female, age 22) were plated at passage 5 at a density of 1000 cells*cm^−2^ in tissue culture flasks and expanded until 80% confluency in complete media (CM) consisting of αMEM with Glutamax and no nucleosides (Gibco) supplemented with 10% fetal bovine serum (FBS) at 37 °C / 5% CO_2_. For the first seeding, hMSCs were incubated for 30 min with CellTracker™ Green (Thermofisher) (5 μm in CM without FBS), rinsed with Dulbecco’s phosphate-buffered saline (dPBS), trypinized and resuspended in CM with 100 U/ml PenStrep (Thermofisher), at a concentration of 4 million cells/ml. A volume of 170 μl of cell suspension was used to seed the scaffolds that were previously disinfected with 70% ethanol for 20 min, rinsed in dPBS and dried inside a laminar flow cabinet on sterile filter paper. Cells were allowed to attach for 4 h at 37 °C / 5% CO_2_. After this time, scaffolds were transferred to a well with 3 ml CM with PenStrep and cultured overnight. For the second seeding, hMSCs were incubated for 30 min with CellTracker™ Deep Red (Thermofisher) (1 μm in CM without FBS), washed, trypinized and resuspended in CM with PenStrep supplemented with 10% dextran (500 kDa, Farmacosmos) at a concentration of 4 million cells/ml. Scaffolds seeded the day before were placed on top of sterile filter paper to remove the media inside the pores and re-seeded with 170 μl viscous cell suspension. Cells were allowed to attach for 4 h and the cultured scaffolds were transferred to a new well with 3 ml CM. After overnight culture, double-seeded scaffolds were fixed using 4% formaldehyde for 20 min, and stored in dPBS until sectioning and imaging. Images were acquired using a fluorescence microscope (Eclipse, Ti2-e, NIKON).

### Statistical analysis

A one-way ANOVA was run on the modulus, strain at failure and strength at failure data using the *anova1* command in MATLAB. Since all results returned p-values < 0.01 for the ANOVA, a Tukey’s post-hoc analysis was then performed using the *multcompare* command in MATLAB.

## General

We thank the Instrument Development, Engineering and Evaluation (IDEE) Department at Maastricht University for their help in manufacturing the printhead. We thank our project partner Gesim for providing the test printer platform, motor controller and assistance with the integration of the printhead and the plasma jet. We thank Andrea Calore for support on the rheology measurements.

We acknowledge the help of Thomas Neubert and Kristina Lachmann (Fraunhofer IST) for initial testing and optimization of plasma polymerization processes; Alberto Sanchez (Tecnalia) for providing the 45nHA material; Marco Scatto (Nadir S.R.L.) for producing the composite materials tested; Erwin Zant (PolyVation B.V.) for providing the polymer; Michele Sisani (Prolabin & Tefarm S.R.L.) for antibiotics fillers; and Rune Wendelbo (Abalonyx) for providing the rGO. Lastly, we thank Prof. Bert van Rietbergen and Giovanni Gonnella (TU Eindhoven) for micro CT imaging of HA gradient scaffolds.

## Funding

This project/research has been made possible with the support of the European Union’s Horizon 2020 research and innovation programme under grant no. 685825 (project website: http://project-fast.eu).

## Author contributions

R.S. carried out the computational modeling of the screw, designed and carried out the validation experiments for the printhead and plasma jet, and prepared the manuscript. M.C.-T. designed and carried out the cell studies, assisted in the design and production of scaffolds, and wrote the relevant parts of the paper. A.P. and P.S. enabled the plasma jet production and wrote relevant parts of the paper. A.P. carried out the plasma jet modeling. E.V.F. assisted with the plasma integration on the printer platform. C.M. led the printhead development and provided the critical design choices. C.M., L.M. and A.P. designed a larger study as part of a consortium project. C.M. and L.M. supervised the work of R.S. and M.C.-T. All authors read and approved the paper.

## Competing interests

P.S. and A.P. own part of the plasma jet manufacturer Nadir s.r.l.. E.V.F. is also affiliated with Nadir. R.S., L.M. and C.M. have applied for a patent (EP18200211.3, submitted: 12 Oct. 2018, PCT/EP2019/077519, submitted: 10 Oct. 2019) on the developed printhead and aim to license it to interested printer companies. M.C.-T. has no financial interests to declare.

