## Supplementary Material for "A hybrid additive manufacturing platform to create bulk and surface composition gradients on scaffolds for tissue regeneration"

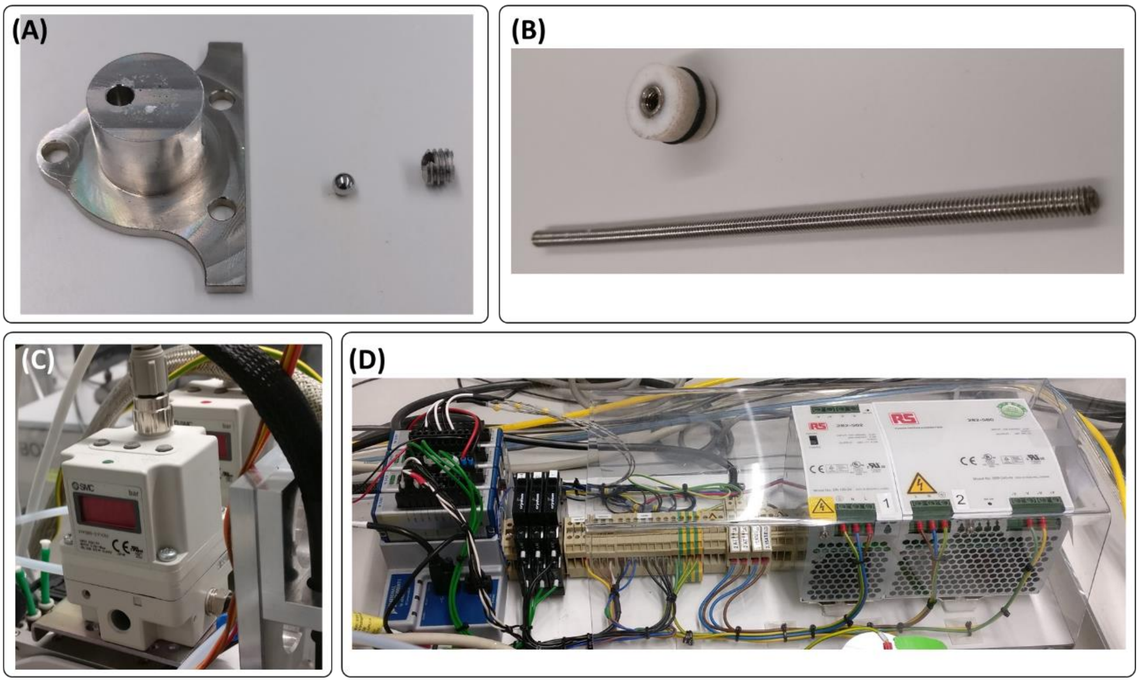


**Supplementary Fig. S1. Several key supporting technologies enable the proper functioning of the printhead.** One-way valves (**A**) at the base of reservoirs prevent back-flows of materials into the other material's reservoir. Extractable plungers (**B**) allow reduction of left-over material in the reservoirs and new material feeding after the reservoirs are emptied. Individual pressure regulators (**C**) and pressure and temperature controllers (**D**) are critical to the printhead operation.

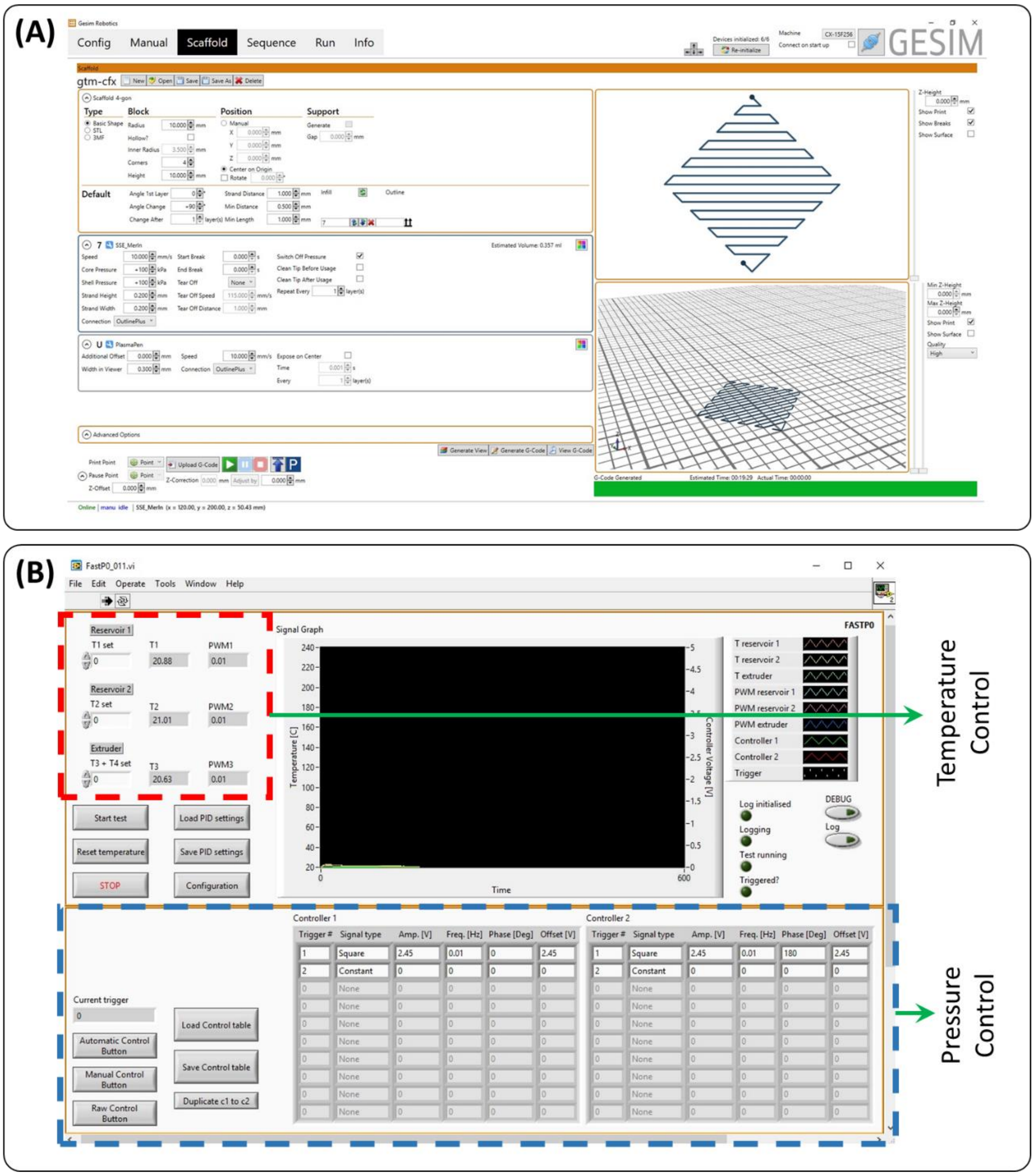


**Supplementary Fig. S2. Customized software enable the application of the desired print control via a user-friendly interface.** The printer software interface (**A**) enables the desired print parameter entry and the automated generation of G-codes to appropriately move and trigger the printhead and the plasma jet. The temperature and pressure software interface (**B**) allows setting the printhead temperatures and the pressure profiles to be applied to each of the printhead reservoirs when triggered.

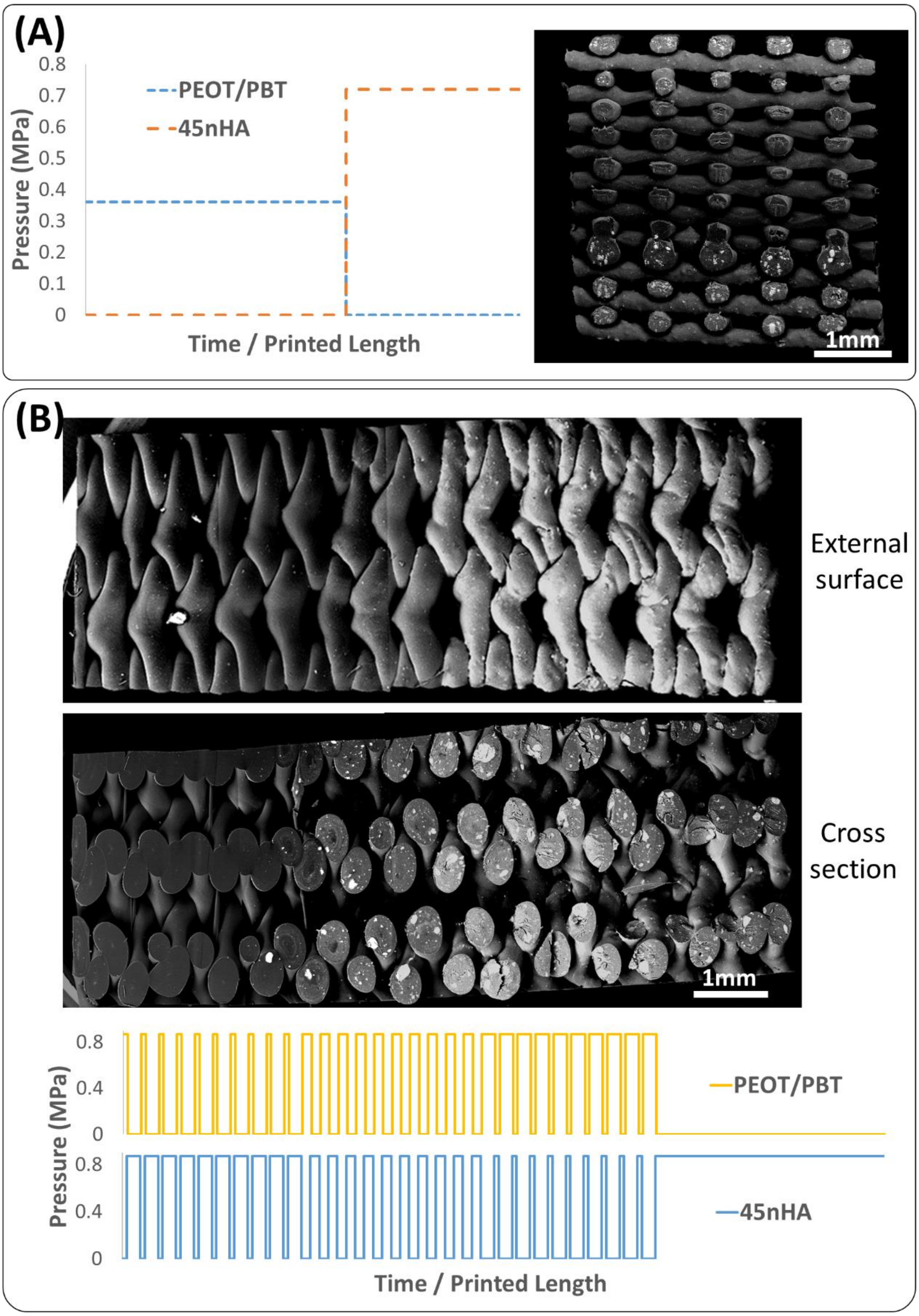


**Supplementary Fig. S3. Continuous complete composition switches as well as sustained intermediate composition printing were demonstrated.** Back-scattered electron-mode scanning electron microscopy (BSE-SEM) images showing HA filler in contrast with the surrounding PEOT/PBT polymer demonstrated that continuous, smooth transitions between compositions could be made when pressures were alternatively applied to the two reservoirs for extended periods of time (**A**). Intermediate compositions could be sustained for a longer duration by alternatively injecting materials into the screw and controlling their ratio by controlling the ratio of times each pressure is on (**B**).

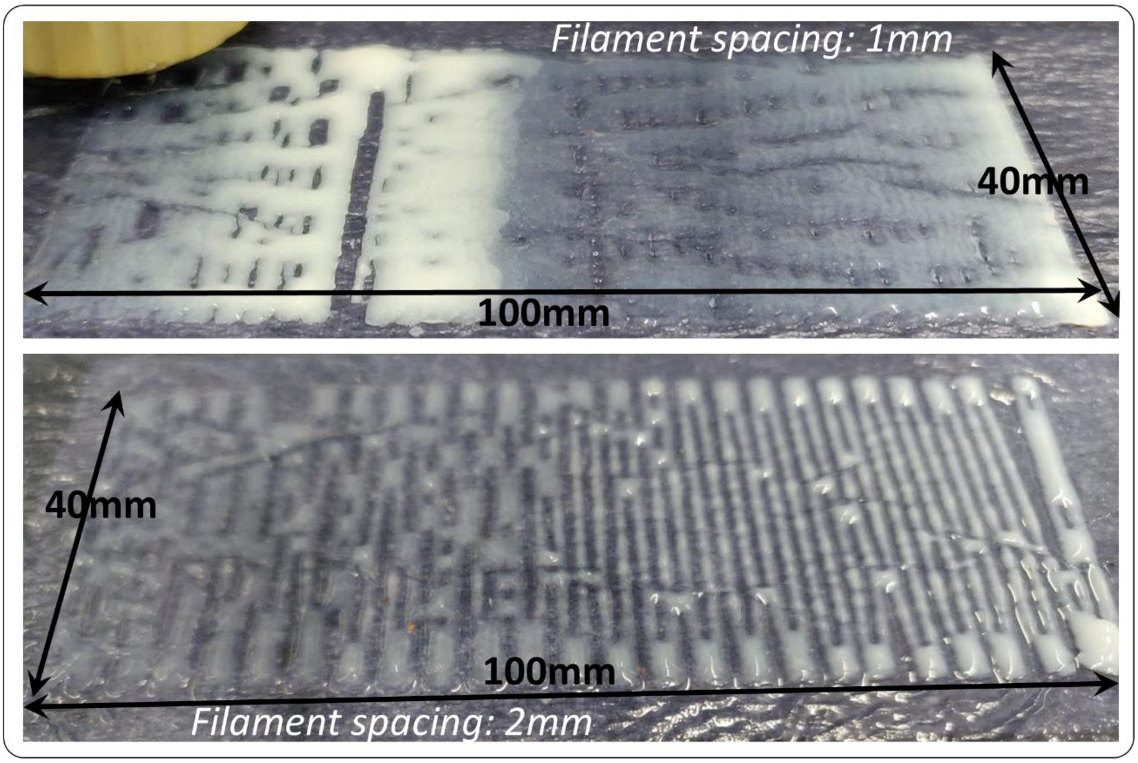


**Supplementary Fig. S4. Continuous composition gradients can be printed using hydrogels.** Alginate gel loaded with hydroxyapatite (HA) was printed on calcium chloride–soaked tissue paper to gel the alginate. Gradients in HA composition could be produced using the printhead.

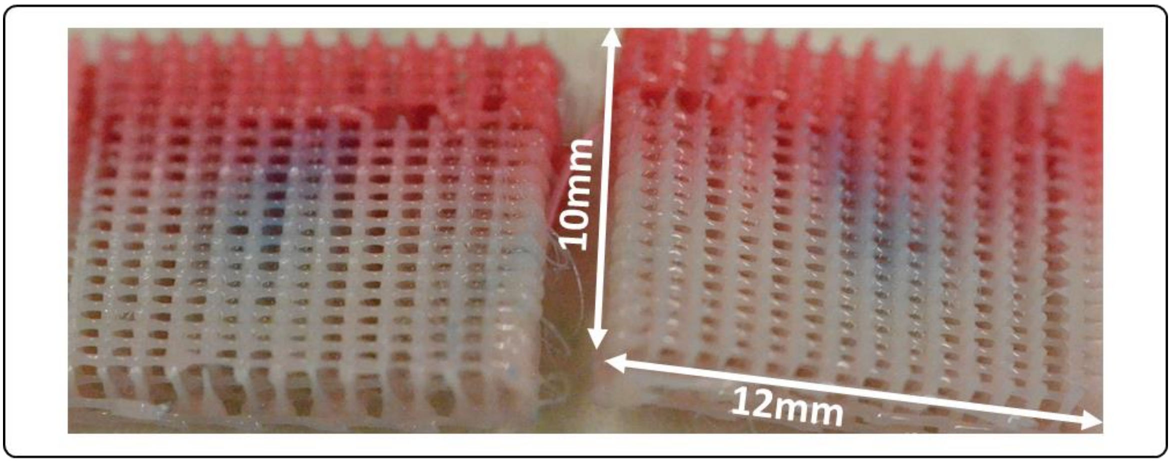


**Supplementary Fig. S5. Bulk composition gradients can be combined with plasma patterning to simultaneously test for the response of cells to various conditions.** Dyed (pink) and non-dyed PEOT/PBT were used to generate the shown composition gradient scaffold, and the core of the scaffold was coated with plasma-polymerized VTMOS-MAA, which is visualized using methylene blue staining.

**Supplementary Table S1. Screw geometrical parameters used for the tested designs in the computational modeling–assisted screw operational torque minimization.** Column values, in mm, correspond to measurements shown in Figure 2A. Design no. 1 has the base dimensions and for the other designs, the parameter changed is highlighted in bold.

| Design No. | t | D1 | D2 | p1 | p2 |
|---|---|---|---|---|---|
| 1 | 1.5 | 4.8 | 5.6 | 8 | 6 |
| 2 | **1** | 4.8 | 5.6 | 8 | 6 |
| 3 | **2** | 4.8 | 5.6 | 8 | 6 |
| 4 | 1.5 | **4.6** | 5.6 | 8 | 6 |
| 5 | 1.5 | **5** | 5.6 | 8 | 6 |
| 6 | 1.5 | 4.8 | **5.4** | 8 | 6 |
| 7 | 1.5 | 4.8 | **5.8** | 8 | 6 |
| 8 | 1.5 | 4.8 | 5.6 | **7** | 6 |
| 9 | 1.5 | 4.8 | 5.6 | **9** | 6 |
| 10 | 1.5 | 4.8 | 5.6 | 8 | **5** |
| 11 | 1.5 | 4.8 | 5.6 | 8 | **7** |

**Supplementary Table S2. Printing parameters for various prints and scaffolds produced.**

| Scaffold type | PEOT/PBT temp. (°C) | PEOT/PBT + filler temp. (°C) | Mixing chamber temp. (°C) | PEOT/PBT pres. profile | PEOT/PBT + filler pres. profile | Screw speed (rpm) | Translation speed (mm/s) | Starting Composition (w/ filler: w/o filler) | Extrusion needle inner diameter (µm) | Strand distance (µm) | Layer height (µm) |
|---|---|---|---|---|---|---|---|---|---|---|---|
| **Empirical switching 45nHA** | 195 | 210 | 220 | 0.88 MPa, 0.01 Hz sq. wave, phase 0 | 0.88 MPa, 0.01 Hz sq. wave, phase 180 | 30 | 6 | 50:50 | 400 | 1000 | 320 |
| **Empirical switching 10rGO** | 195 | 220 | 220 | 0.45 MPa, 0.002 Hz sq. wave, phase 180 | 0.88 MPa, 0.002 Hz sq. wave, phase 0 | 60 | 3 | 50:50 | 250 | 1000 | 200 |
| **Empirical switching 20LDH-CFX** | 195 | 185 | 195 | 0.88 MPa, 0.01 Hz sq. wave, phase 0 | 0.88 MPa, 0.01 Hz sq. wave, phase 180 | 60 | 10 | 50:50 | 400 | 1000 | 320 |
| **Empirical switching 20ZrP-GTM** | 195 | 190 | 195 | 0.88 MPa, 0.005 Hz sq. wave, phase 0 | 0.88 MPa, 0.005 Hz sq. wave, phase 180 | 60 | 6 | 50:50 | 400 | 1000 | 320 |
| **Mech. Test – 45nHA** | 195 | 210 | 220 | 0 | 0.72 | 60 | 22.5 | 100:0 | 250 | 750 | 200 |
| **Mech. Test – PEOT/PBT** | 195 | 210 | 220 | 0.36 | 0 | 60 | 17.5 | 0:100 | 250 | 750 | 200 |
| **Mech. Test – HPH_C** | 195 | 210 | 220 | 0.36 MPa for layers 1–13 | 0.72 MPa for layers 14–20 | 60 | 17.5 for first 15 layers, 22.5 for next 5 layers | 100:0 | 250 | 750 | 200 |
| **Mech. Test – HPH_D** | 195 | 210 | 220 | 0.36 MPa for layers 1–5 and 16–20 | 0.72 MPa for layers 6–15 | 60 | 17.5 for first 5 and last 5 layers, 22.5 for middle 10 layers | 100:0 | 250 | 750 | 200 |
| **Mech. Test – PHP_C** | 195 | 210 | 220 | 0.54 MPa for layers 1–3 and 14–32 | 0.72 MPa for layers 4–13 | 20, varied manually a few times | 7.5 for layers 15–20, 15 for rest of the layers | 0:100 | 340 | 850 | 250 |
| **Mech. Test – PHP_D** | 195 | 210 | 220 | 0.54 MPa for layers 1–11 and 22–32 | 0.72 MPa for layers 12–21 | 20 | 15 | 0:100 | 340 | 850 | 250 |

**Movie S1. Comparison of particle mixing between screws with a triangular and a square thread cut.**

**Movie S2. Segmental long bone defect scaffold printing with a cortical composition change.**

**Movie S3. Sustained printing of compositions intermediate between the compositions in the two reservoirs.**

**Movie S4. Mechanical testing of continuous and discrete gradient scaffolds with a PEOT/PBT region sandwiched between 45nHA regions.**

**Movie S5. Mechanical testing of continuous and discrete gradient scaffolds with a 45nHA region sandwiched between PEOT/PBT regions.**

**Movie S6. Scaffold production combining a continuous bulk composition gradient and plasma patterning.**
